# Impact of integration on persistent homology clustering and biological signal detection in scRNA-seq data

**DOI:** 10.1101/2025.07.24.666637

**Authors:** Jonah Daneshmand, Julia H. Chariker, Akshitkumar Mistry, Eric C. Rouchka

## Abstract

**Background:** As the availability of single-cell RNA sequencing (scRNA-seq) data expands, there is a growing need for robust methods that enable integration and comparison across diverse biological conditions and experimental protocols. Persistent homology (PH), a technique from topological data analysis (TDA), provides a deformation-invariant framework for capturing structural patterns in high-dimensional data.

**Methods:** In this study, PH was applied to a diverse collection of scRNA-seq datasets spanning eight tissue types to investigate how data integration affects the topological features and biological interpretability of the resulting representations. Clustering was performed based on PH-derived pairwise distances and global topological structure was assessed through Betti curves, Euler characteristics, and persistence landscapes. By comparing these summaries across raw, normalized, and integrated datasets, we examined whether integration enhances the detection of biologically meaningful patterns, or, conversely, obscures fine-scale structure.

**Results:** This approach demonstrates that PH can serve as a powerful complementary strategy for evaluating the impact of integration and reveals how topological summaries can help disentangle biological signal from batch-related noise in single-cell data. This work establishes a framework for using topological methods to assess integration quality and highlights new avenues for interpreting complex transcriptomic landscapes beyond conventional clustering.

## INTRODUCTION

Single-cell RNA-sequencing (scRNA-seq) has revolutionized cellular transcriptomics by enabling detailed gene expression analysis at a single cell resolution [1]. This level of granularity allows comprehensive characterization of cellular heterogeneity, interactions, and states across various biological contexts [2]. With rapid growth in publicly available scRNA-seq datasets, effective methods for integrating and comparing data have become crucial [3].

ScRNA-seq data exhibit inherent sparsity and zero-inflation due to dropout events, in which expressed genes remain undetected during sequencing [4]. This noise and variability, driven by biological stochasticity and technical factors such as sequencing depth and library preparation, can severely impact downstream analyses, leading to misclassification during clustering, difficulty in identifying rare cell populations, and reduced sensitivity in differential expression studies [5].

Computational integration tools, including Seurat [6], Harmony [7], LIGER [8], and Scanorama [9], employ strategies for both integration and harmonization. Integration involves merging datasets into a unified representation, whereas harmonization specifically targets the removal of technical variability without distorting underlying biological signals [10]. Methods such as canonical correlation analysis (CCA) [11], reciprocal principal component analysis (RPCA) [11], iterative clustering [12], integrative non-negative matrix factorization (iNMF) [8], mutual nearest neighbor alignment [13], and Harmony [7] can mitigate technical biases but differ in their ability to preserve rare cell populations or subtle transcriptional states [14]. Despite their utility, integration methods risk obscuring biologically critical variations. Subtle transcriptional differences can be inadvertently masked if they overlap significantly with batch-specific effects [14]. Consequently, systematic evaluation of these integration approaches is essential to ensure biologically meaningful patterns are enhanced rather than diminished.

Persistent homology (PH) and topological data analysis (TDA) offer robust analytical frameworks that focus on relational and structural features of datasets, notably the branching architectures relevant to cellular differentiation trajectories often examined in scRNA-seq [15–17]. PH has been applied to structural biology [18–19], gene expression patterns in tumors [20], viral phylogenetics [21], and cancer cellular architectures [22]. In scRNA-seq contexts, PH and TDA have revealed patterns in cellular differentiation and gene co-expression networks, even under substantial noise [23–24].

### Challenges in integrating and analyzing diverse datasets

Integrating multi-tissue scRNA-seq datasets presents several key challenges to unified downstream analysis. First, the intrinsic transcriptional heterogeneity across tissues can dominate integrated representations, making it difficult to retain subtle, tissue-specific cell states. Second, both overlapping and divergent cell populations introduce harmonization difficulties [14]. Third, variation in biological context adds another layer of heterogeneity that, without careful modeling, risks being conflated with technical noise. Fourth, technical batch effects stemming from differences in sequencing platforms, library preparation methods, and laboratory protocols can obscure true biological variation [25–26]. Fifth, disparities in data sparsity across tissues can bias clustering results and downstream differential expression analyses [27–28].

Taken together, these challenges highlight the difficulty of extracting consistent, biologically meaningful structure from integrated datasets. PH offers a complementary strategy by quantifying higher-order topological features robust to deformation and capable of capturing global organizational patterns, even in the presence of batch effects and non-uniform sampling. By applying PH across raw and integrated datasets, we aim to assess the degree to which topological methods preserve or recover biological structure amid these integration challenges.

### PH and its topological basis

PH is a technique from TDA that leverages deformation-invariant features to mitigate batch effects and technical noise in scRNA-seq studies [27]. PH robustly identifies biological patterns and improves interpretability in integrated analyses by quantifying persistent structural elements (connected components, loops, and voids) through their birth and death thresholds across filtration scales (Figure 1) [1, 30–34].

**Figure 1:**
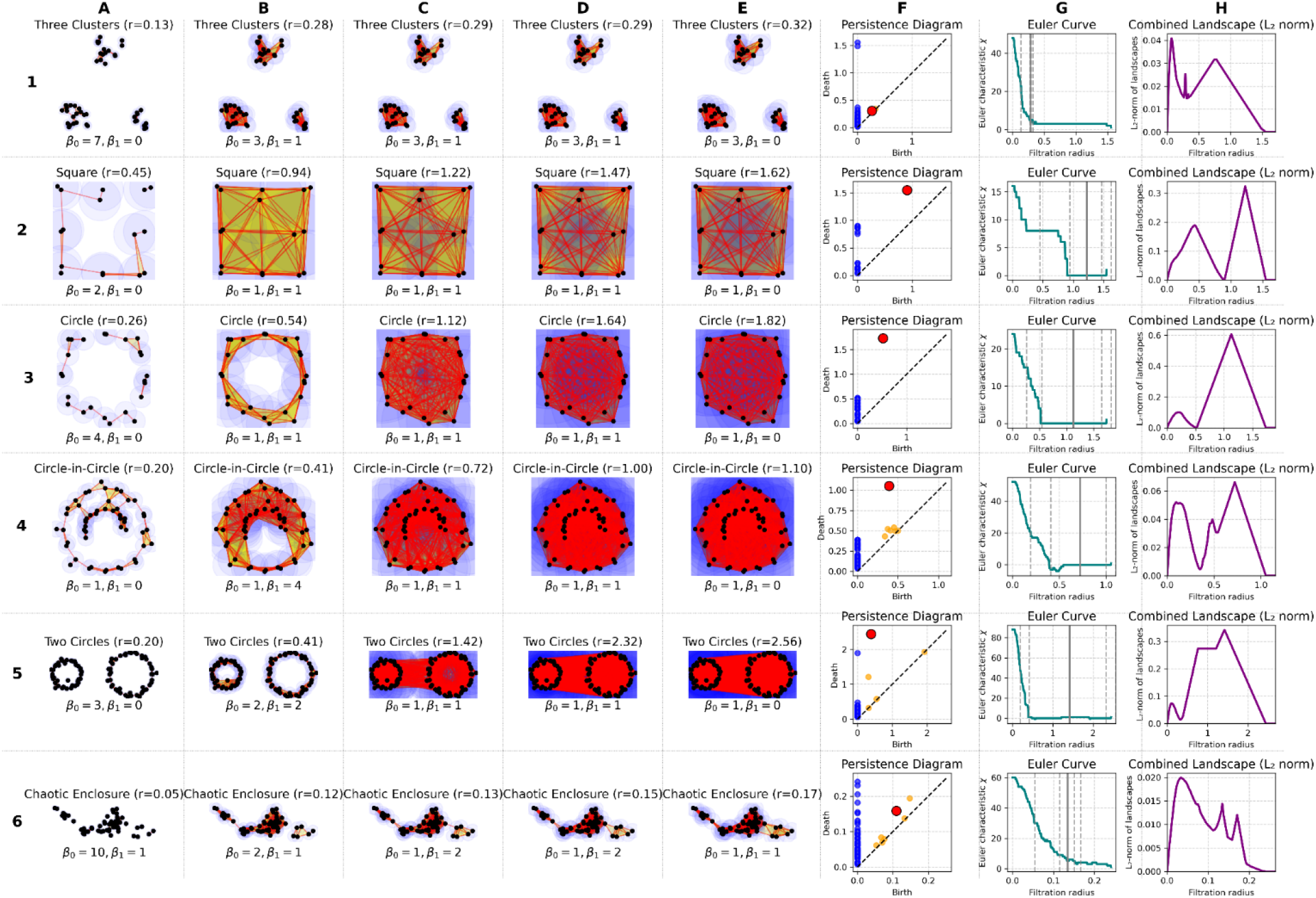
Topological analysis of six synthetic point clouds using Vietoris–Rips filtration and PH. The TDA pipeline is applied to six datasets, with each row analyzing a distinct shape and each column a different analytical step. Rows analyze six datasets: (1) Three Clusters, (2) Square, (3) Circle, (4) Circle-in-Circle, (5) Two Circles, and (6) a Chaotic Enclosure. (A-E) Snapshots of the Vietoris-Rips filtration at key radii. These radii are dynamically selected to show the lifecycle of the most persistent 1-dimensional hole (H_1_): (B) just after birth, (C) mid-life, (D) just before death, and (E) after the hole is filled. (F) The Persistence Diagram, summarizing the birth and death of connected components (H_0_) and holes (H_1_) over the entire filtration. (G) The Euler Characteristic Curve, a 1-D topological summary calculated as the number of connected components minus the number of holes (χ=β_0_−β_1_). (H) The L_2_-norm of the persistence landscape, a stable vector representation suitable for machine learning. This process systematically converts unstructured point data into quantitative topological signatures.

A hallmark of PH is its resilience to small, continuous distortions (e.g., a global shift in gene expression values caused by batch effects) that do not alter the fundamental shape of the point cloud. Consequently, core topological features such as connected components or loops remain intact across datasets. Emphasizing these relational structures, rather than absolute expression values, significantly reduces noise sensitivity and enhances interpretability, ultimately reinforcing PH’s utility for scRNA-seq integration.

### Evaluation of integration methods in PH-based clustering

Evaluating integration methods through PH-based clustering provides insight into their effectiveness in preserving biological signals within scRNA-seq data. PH captures topological features independently of linearity assumptions or data distribution normality, offering advantages over traditional evaluation metrics [33]. By comparing integrated and non-integrated topologies, PH reveals how harmonization strategies impact biologically meaningful clustering [34]. We sought to define the conditions under which PH provides the most biologically relevant insights across raw, normalized, and integrated single-cell datasets.

## MATERIAL AND METHODS

### Datasets

Publicly available scRNA-seq datasets from the Gene Expression Omnibus (GEO) available as RData files from PanglaoDB [35] were utilized. The original data are linked via their Sequence Read Archive (SRA) identifiers, ensuring traceability of data provenance. Comprehensive details are provided in Supplemental Table 1 and Supplemental Table 2. The primary dataset (Supplemental Table 1) encompasses eight human tissues: substantia nigra, pancreatic islets, bone marrow, liver, testis, colon, prostate, and peripheral blood mononuclear cells (PBMCs). All samples in this primary collection were sequenced using the 10x Genomics Chromium platform, ensuring consistency in the underlying sequencing technology. For validation purposes, a smaller, homogeneous dataset (GSE120221; Supplemental Table 2), consisting of bone marrow samples was used.

Initial data preparation involved loading the RData files into Seurat objects and embedding detailed metadata, including SRA identifiers, tissue type, and sequencing approach (scRNA-seq or single-nucleus RNA-seq). Standard quality control (QC) was then applied: cells were filtered if they expressed fewer than 500 genes, representing cell-free RNA, or more than 9,000 genes, representing potential doublets. Cells were also removed if they exhibited greater than 20% mitochondrial gene content or less than 5% ribosomal gene expression. Genes detected in fewer than three cells were removed. All datasets were systematically verified for metadata consistency and completeness.

### Computational approaches

All analyses were implemented in R (v4.4.1) with core libraries loaded at the outset: tidyverse (v2.0.0), Matrix (v1.7-3), Seurat (v5.3.0), SeuratObject (v5.0.2), SeuratDisk (v0.0.0.9021), ripserr (v0.3.0), TDA (v1.9.4), TDAstats (v0.4.1), foreach (v1.5.2), doParallel (v1.0.17), parallel (4.4.1), kernlab (v0.9-33), igraph (v2.1.4), ComplexHeatmap (v2.20.0), circlize (v0.4.16), viridis (v0.6.5), dendextend (v1.19.0), mclust (v6.1.1) and aricode (v1.0.3), plus the utility packages plyr (v1.8.9), stringr (v1.5.1), reshape2 (v1.4.4), gridExtra (v2.3), RColorBrewer (v1.1-3), digest (v0.6.37), transport (v0.15-4), scCustomize (v3.0.1), BiocSingular (v1.20.0), Rtsne (v0.17), and patchwork (v1.3.0). Additional dependencies and version numbers are provided in Supplemental Table 3. Our analysis functions and pipeline are available on GitHub (https://github.com/jcdaneshmand/scPHcompare).

A comprehensive analytical workflow was implemented to evaluate the impact of integration on the topological structure of scRNA-seq data. The process began with the acquisition and preprocessing of data (described above) for both the comprehensive and test datasets. Raw count matrices were imported into Seurat (v5.2.1) objects using Read10X_h5 or custom loadRData functions.

Following initial preprocessing, four distinct normalization and integration variations were generated for comparative analysis: (1) the original unnormalized (Raw) data; (2) data normalized with SCTransform [36] (SCT Individual); (3) datasets merged then collectively normalized with SCTransform (SCT Whole); and (4) harmonization using Seurat’s integration workflow following SCT Individual (Integrated).

The core of the analysis involved PH computation. For each data iteration and sample, pairwise cell-distance matrices were constructed and Vietoris–Rips filtration was applied to generate persistence diagrams (PDs) using ripserr (v0.3.0). These PDs captured the birth and death profiles of essential topological features, such as connected components (β_0_) and loops (β_1_). To facilitate quantitative comparisons and visualization of topological differences, these PDs were subsequently converted into summary statistics, including Betti curves, Euler curves, and persistence landscapes.

Pairwise statistical comparisons were conducted to quantify differences in topological signals across known biological groups. Specifically, Euler curves (calculated as β_0_ −β_1_) and aggregated persistence landscapes (formed by combining β_0_ and β_1_ information via an L^2^ norm) were compared. The statistical significance of observed differences was evaluated at each filtration threshold using Kolmogorov–Smirnov (KS) tests, alongside bootstrap and permutation-based strategies, with false-discovery rate (FDR—Benjamini-Hochberg) corrections applied to p-values. These pointwise tests were summarized by identifying minimum FDR and maximum KS statistics, and global permutation tests were performed over the entire filtration range for an overall assessment.

In parallel, topological-based hierarchical clustering and statistical evaluation was performed. Multiple topological distance matrices from the PDs and landscapes were derived, including bottleneck distance, spectral distance, and landscape distance matrices. Both hierarchical and spectral clustering were applied to assess their ability to group samples according to known labels. The statistical significance of the resulting clustering performance was quantified using a bootstrap null-model, comparing against standard metrics including Adjusted Rand Index (ARI), Normalized Mutual Information (NMI), Jaccard Index, Variation of Information (VI), and cluster purity.

Standard clustering was performed using Seurat’s Louvain algorithm and K-means clustering on principal components for each of the four data representations (Raw, SCT Individual, SCT Whole, Integrated). The performance of these conventional methods was directly compared to the topological clustering results to evaluate preservation of biologically relevant groupings.

How data integration procedures can enhance or, conversely, obscure the detection of biologically relevant topological features within scRNA-seq data was elucidated by juxtaposing topological summaries (Betti, Euler, and landscape curves) and clustering metrics across these multiple normalization and integration pipelines.

### Normalization and integration variations

Four distinct data representations (Raw, SCT Individual, SCT Whole, and Integrated) were generated from the QC count matrices to evaluate how preprocessing influences the topological features captured by PH.

#### Raw

The Raw representation consists of the filtered raw count data for each sample, with no additional normalization applied. This serves as a baseline condition, preserving the original signal and technical noise, against which all other processed iterations are compared.

#### SCTransform Individual (SCT Individual)

In the SCT Individual representation, each sample’s filtered raw count matrix was independently normalized using SCTransform (SCT) [36]. The step of regressing out technical covariates, such as mitochondrial and ribosomal gene percentage, was skipped to preserve technical variation for topological assessment. This allows for evaluating PH’s capacity to extract biological structure without prior removal of known technical artifacts.

#### SCT Whole

SCT Whole was generated by merging all individually filtered raw count matrices into a single dataset, followed by joint normalization using SCTransform. As with the SCT Individual representation, technical variable regression was omitted. This allows for evaluating how joint normalization without batch correction or technical regression affects topological structures, particularly compared to individually normalized data.

#### Integrated

The Integrated representation was derived by applying Seurat’s integration workflow to the objects produced in the SCT Individual step. Integration proceeded as follows: First, shared variable features were identified using SelectIntegrationFeatures (default n=3000). Then PrepSCTIntegration prepared the data using these features. Anchors were computed with FindIntegrationAnchors, using SCT normalization and reciprocal principal component analysis (RPCA) for dimensional reduction. RPCA allows for scalability and robustness, especially when integrating datasets with varying cell numbers or complex batch effects. If RPCA failed, canonical correlation analysis (CCA) was used as a fallback. Integration was completed using IntegrateData, which utilized the identified anchors and the k.weight parameter (default 50) to harmonize the datasets into a shared latent space, producing the final integrated assay. As in previous iterations, no technical variable regression was performed.

### PH workflow

PH computation was applied to the expression matrices derived from each sample within the four preprocessing iterations. For each sample, the input for PH consisted of its cell-by-gene expression matrix (using either raw counts or normalized/integrated values, depending on the iteration). Vietoris-Rips filtration was then performed. While Euclidean distance was employed, alternative metrics (e.g., correlation-based) could offer complementary insights and were considered for future exploration.

This filtration process yielded PDs for each sample, which summarize the birth and death scales of topological features, primarily connected components (dimension 0, β_0_) and loops (dimension 1, β_1_), across the filtration range. To facilitate quantitative comparison and interpretation, these PDs were transformed into several summary statistics.

Betti curves were computed from PDs separately for each dimension (β_0_(t) and β_1_(t)), quantifying the number of persistent topological features present at each filtration threshold t. Euler curves (χ(t)) were derived by integrating information across dimensions (χ(t)=β_0_(t)−β_1_(t)), providing a holistic summary of topological complexity as a function of the filtration scale. These curve-based descriptors summarize key topological characteristics, facilitating robust quantitative comparisons of structural variations across different datasets, preprocessing iterations, and biological groups (e.g., tissues) to assess the biological relevance of observed differences.

Persistence landscapes were computed from the PDs. Landscapes offer an alternative topological summary by transforming the discrete PD points into a stable functional representation suitable for statistical analysis and clustering. Specifically, each feature (birth-death pair) in a PD is mapped to a “tent” function in the landscape space; the landscape function at a given level k and filtration value t is the kth largest value among all tent functions at t. This representation captures feature prominence and persistence across scales. Persistence landscapes reside in a Banach space, which allows for direct computation of means, variances, and other statistics on the topological summaries and facilitating the application of standard statistical techniques, while also offering improved numerical stability compared to PDs.

Persistence landscapes were computed separately for dimension 0 and dimension 1 from each PD. Specifically, the first landscape function (k=1), which captures the most prominent topological features, was evaluated over a discretized grid of 100 equally spaced points along the filtration scale for both dimension 0 (L_0_(t)) and dimension 1 (L_1_(t)). To streamline downstream analysis, these first-level landscape functions were combined into a single scalar-valued aggregated landscape curve for each sample. This aggregation was performed pointwise across the discretized filtration scale. Specifically, for each grid point t_i_ (where i=1,…,100), the aggregated landscape value L_agg_(t_i_) was computed as the L_2_ (Euclidean) norm of the corresponding first-level landscape values from dimension 0 (L_0_(t_i_)) and dimension 1 (L_1_(t_i_)): L_agg_(t_i_)=(L_0_(t_i_))^2^+(L_1_(t_i_))^2^.

### Clustering analysis

Clustering analyses were performed using both topologically derived distances and conventional clustering approaches to evaluate the preservation of biological structure across different preprocessing strategies. Three distinct pairwise distance matrices were derived from their respective PDs and persistence landscapes to quantify dissimilarities between the topological signatures of different samples. First, Bottleneck Distance Matrices (BDMs) were constructed directly from the PDs of each sample. The bottleneck distance was computed between the PDs of every pair of samples. This distance measures the cost of the optimal matching between features in two diagrams, considering both dimension 0 (connected components) and dimension 1 (loops). The distance reported is the maximum of the bottleneck distances calculated for each dimension separately. Second, Spectral Distance Matrices (SDMs) were derived from the BDMs to capture inter-sample dissimilarity. This involved converting the BDM into a similarity matrix using a Radial Basis Function (RBF, Gaussian) kernel, with the kernel width (σ) adaptively scaled based on the BDM’s range. The normalized graph Laplacian was then computed from this similarity matrix. Spectral decomposition of the Laplacian provided an embedding of the samples into a lower-dimensional space defined by the eigenvectors corresponding to the smallest non-trivial eigenvalues (typically the first 50, excluding the trivial one). The SDM was then calculated as the matrix of pairwise Euclidean distances between samples in this new spectral embedding. Third, Landscape Distance Matrices (LDMs) were generated from the persistence landscapes. For each sample, the first landscape function (L_0_(t) for dimension 0 and L_1_(t) for dimension 1, evaluated on a common grid) was used. The pairwise distance in the LDM between any two samples i and j was computed by first determining the L_2_ (Euclidean) distance between their respective dimension 0 landscapes (d_0_=∣∣L_0,i_−L_0,j_∣∣^2^) and dimension 1 landscapes (d_1_ =∣∣L_1,i_−L_1,j_∣∣^2^). The final LDM entry (i,j) was the Euclidean combination of these distances: LDM_i,j_=d_02_+d_12_. This approach provides a distance metric based on the functional representation of topological features, offering a complementary perspective to the PD-based BDM and SDM. Collectively, these three matrices (BDM, SDM, and LDM) provided complementary measures of topological dissimilarity between samples. These matrices served as inputs for both hierarchical clustering (using Ward’s linkage) and spectral clustering. The number of clusters (k) for these methods was predefined based on known biological or experimental attributes, such as tissue type, SRA study, run accession, or sequencing approach. In addition, randomized groupings were used as a negative control. These biologically anchored cluster assignments allowed us to evaluate how well the topological patterns captured by different distance metrics and clustering algorithms aligned with known sample characteristics, thereby assessing the biological relevance of the detected topological variations.

To complement the topologically driven clustering, conventional clustering methods were applied. These included Seurat-based graph clustering (Louvain algorithm) and K-means clustering performed on the principal components derived from each data iteration (Raw, SCT Individual, SCT Whole, Integrated). These conventional methods were executed independently on each of the four data representations, generating cluster assignments at biologically relevant resolutions (e.g., based on the number of known tissue types).

### TDA and clustering statistical evaluation with bootstrap strategies

To ensure robust validation, a null-model bootstrap strategy was incorporated to evaluate the statistical significance of clustering performance by comparing observed metric values against those obtained from randomized label assignments. An analogous approach was applied for Betti, Euler, and Landscape curves, where empirical null distributions for group-average curves were generated by permuting sample labels between the compared groups. This involved randomly reassigning the actual sample-derived persistence diagrams (and their corresponding Betti/Euler curves) to the two groups being tested, then recalculating group-average curves for this permuted assignment. Similarly, for aggregated persistence landscapes (where dimensions β_0_ and β_1_ were combined into a single curve per sample using the L_2_ norm at each filtration threshold), sample-level aggregated landscape curves were randomly reassigned to synthetic groups to establish empirical null distributions.

At each filtration threshold (τ) along these curves, KS tests were used to evaluate the differences between the distributions of curve values from two groups. These pointwise tests were summarized by identifying the minimum FDR and the maximum KS statistic across all τ values. Additionally, global permutation tests were performed by integrating the differences between mean curves over the entire filtration range and comparing this integrated statistic to a null distribution generated by permuting sample labels between groups. This unified analytical framework quantified the statistical significance of topological differences consistently across Betti curves, Euler curves, and aggregated landscape curves.

Within-group variability for Betti, Euler, and aggregated landscape curves was assessed by bootstrapping at the sample level. For each biological group, individual sample curves were resampled with replacement to generate mean curves and confidence intervals.

A null distribution framework was also applied to evaluate clustering performance. For each clustering method tested, a null distribution of clustering metric values (ARI, NMI) was generated by repeatedly computing these metrics on data with randomly permuted group labels. This allowed for the estimation of empirical p-values and normalized effect sizes, helping to identify clustering outcomes that significantly outperformed random expectation. This evaluation framework was applied uniformly across all topological distance matrices and conventional clustering methods, allowing for direct comparisons of performance. Among these, ARI and NMI were prioritized for interpretation due to their robustness to varying cluster sizes and clear interpretability when ground truth labels are available.

Cross-iteration robustness of biological signals was evaluated by comparing both the topological summary curves (Betti, Euler, aggregated landscape) and the clustering outcomes (using ARI/NMI) for specific biological groupings across the different preprocessing strategies. For clustering comparisons across iterations where cluster identities might shift, methods such as matching clusters based on the proximity of their centroids in a common principal component space (derived from the integrated data) were considered to align clusters before computing similarity, although primary evaluation focused on the ability to recover known ground-truth labels. Statistical significance of differences in performance was assessed using bootstrap-permuted null models where appropriate.

## RESULTS

### Clustering performance across integration methods

#### Smaller validation dataset collection (bone marrow; GSE120221)

Clustering performance was evaluated for the validation set against the SRA accession (individual sample ID) as the ground truth, with k=25 (Figure 2). As a foundational validation, PH-based distance metrics were tested to see if they could resolve individual samples in a controlled setting. When the number of clusters is set to the number of samples (k=25), hierarchical clustering is expected to achieve a perfect score provided the underlying distance matrix assigns a unique, non-zero distance between distinct samples. Indeed, across all four data iterations, hierarchical clustering based on all three PH-derived metrics yielded perfect performance, with Normalized Adjusted Rand Index (ARI_Norm) and Normalized Mutual Information (NMI_Norm) scores of 1.0 (Supplemental Table 4).

**Figure 2:**
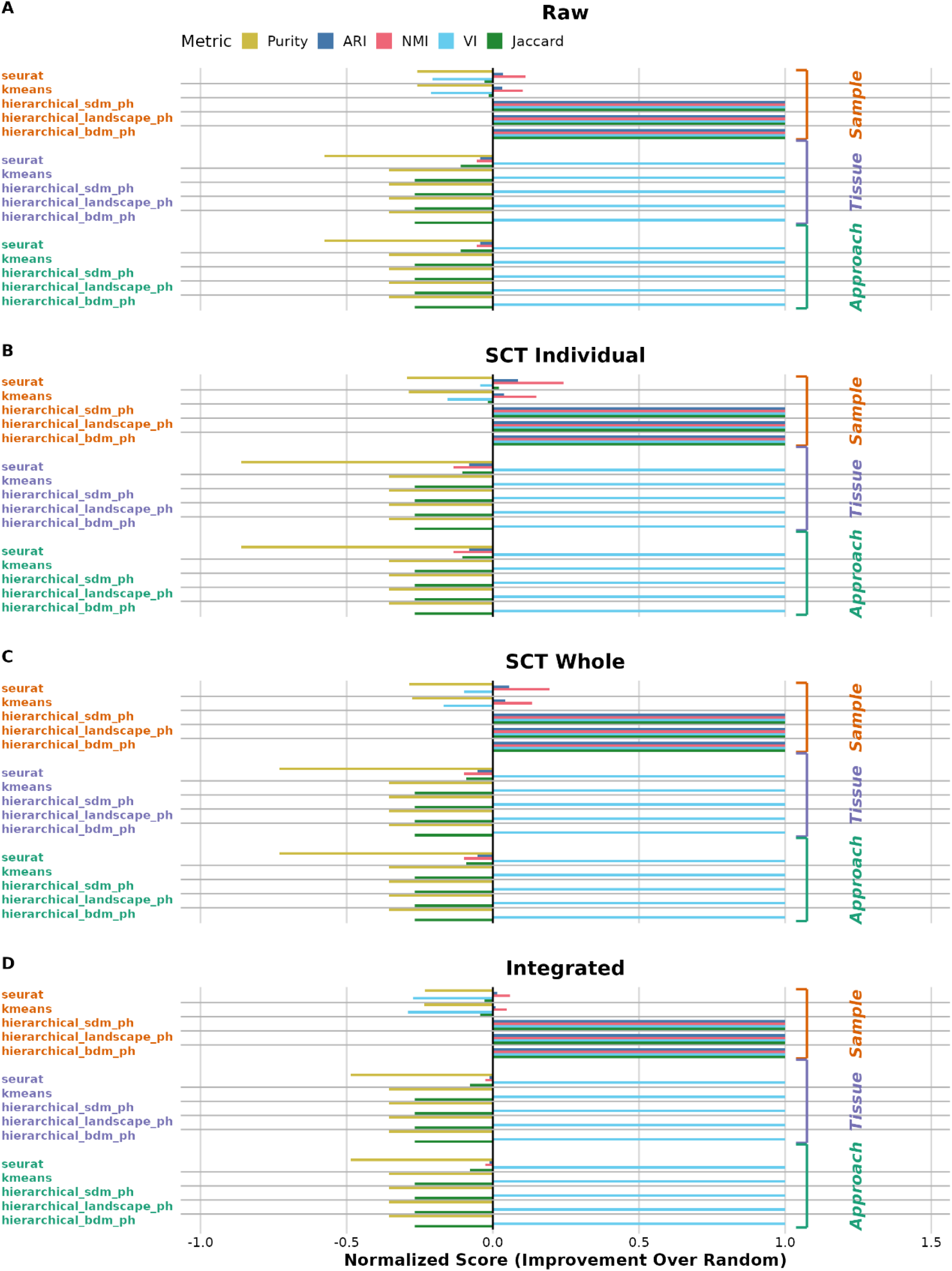
Normalized Cluster Performance Metrics on the Bone Marrow Dataset Collection across Iterations. The figure displays a comprehensive evaluation of clustering performance on the homogeneous bone marrow dataset (GSE12022). Each panel represents a distinct data preprocessing iteration: (A) Raw, (B) SCT Individual, (C) SCT Whole, and (D) Integrated. For each panel, clustering methods (y-axis) are grouped by the reference label they were evaluated against (Sample, Tissue, or Approach). Performance is quantified using five clustering metrics, indicated by color: Adjusted Rand Index (ARI), Normalized Mutual Information (NMI), Jaccard Index, Purity, and Variation of Information (VI). Each “Observed” metric value is a single global score calculated by comparing the full set of predicted cluster labels against the true reference labels for all cells. Because this dataset consists of a single tissue type from a single sequencing approach, the ‘Tissue’ and ‘Approach’ comparisons are degenerate cases where all cells have the same true label, making the performance metric uninformative. The x-axis shows the Normalized Score, which represents the improvement of a method’s performance over a random baseline. This score is calculated by taking the observed metric value, subtracting the mean value from a set of random clustering permutations, and then scaling the result. For metrics where a higher value is better (ARI, NMI, Jaccard, Purity), the formula is (Observed - Random Mean) / (1 - Random Mean). For Variation of Information (VI), where lower is better, the formula is (Random Mean - Observed) / Random Mean. A score of 1 indicates the best possible performance relative to the random baseline, a score of 0 indicates performance equivalent to random, and negative scores indicate performance worse than random. Spectral clustering methods were excluded from this validation analysis as they are not applicable for k=1 (for the single tissue type and approach) or k=25 (for the number of SRA samples), as the algorithm fails when the underlying graph is fully connected.

Conventional clustering methods, K-means and Seurat Louvain, also successfully distinguished individual samples, outperforming random chance (all associated p-values for ARI and NMI were 0 after FDR correction). Across all iterations, K-means yielded ARI_Norm values ranging from 0.0100 to 0.0432 and NMI_Norm values from 0.0481 to 0.1491. Similarly, Seurat Louvain clustering produced ARI_Norm values between 0.0151 and 0.0861 and NMI_Norm values from 0.0588 to 0.2422 (Supplemental Table 4).

While all these ARI_Norm and NMI_Norm values for K-means and Seurat Louvain clustering are positive, indicating performance better than random, the scores are modest. Notably, these scores did not substantially differ across the Raw, SCT Individual, SCT Whole, and Integrated data iterations (Figure 2). This suggests that for the test dataset, neither individual normalization (SCT Individual), collective normalization (SCT Whole), nor full integration (Integrated) offered substantial additional benefits in resolving these known sample identities beyond what was achievable with raw data using these methods.

Evaluations using other potential reference labels, such as tissue type or sequencing approach, were effectively uninformative in this single-tissue, single-platform scenario. As expected, clustering against these uninformative labels resulted in ARI_Norm and NMI_Norm values at or below zero, confirming they did not provide further comparative insights for clustering method efficacy.

The perfect scores confirm that all three PH-based distance calculations successfully generate distinct topological signatures for non-identical samples, thereby establishing technical integrity of our pipeline. With this foundational capability confirmed, we evaluated how these methods perform in a heterogeneous, multi-tissue dataset.

#### Heterogeneous dataset collection (eight human tissues)

Clustering performance on the primary dataset comprised of eight distinct human tissues was assessed to evaluate how well different clustering methods could recover known biological groupings across the data preprocessing iterations. Normalized clustering performance metrics for various topological and conventional methods are presented in Figure 3, with detailed values available in Supplemental Table 5.

**Figure 3:**
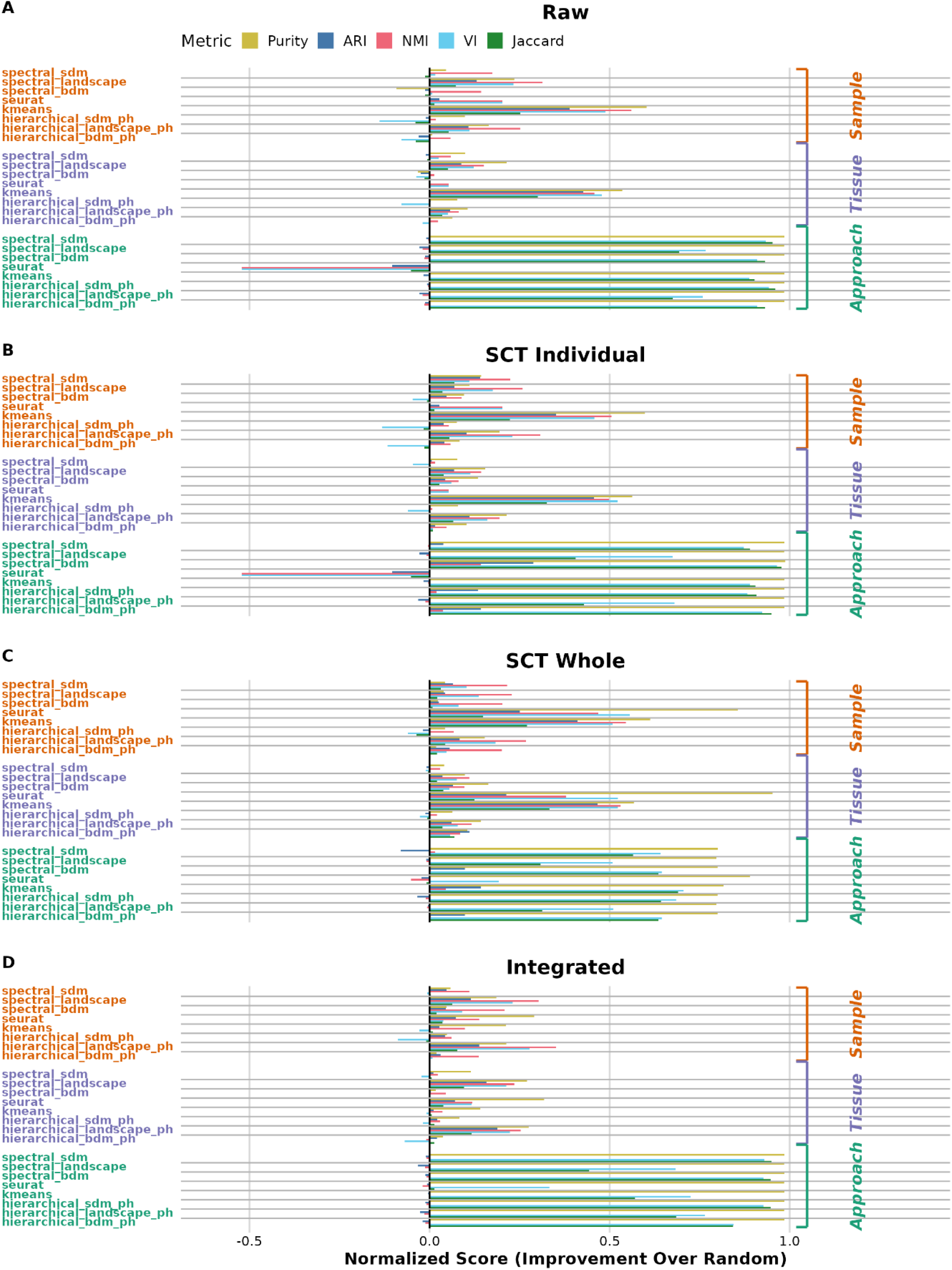
Normalized Cluster Performance Metrics on the Multi-Tissue Dataset Collection across Iterations. The figure displays a comprehensive evaluation of clustering performance on the heterogeneous eight-tissue dataset. Each panel represents a distinct data preprocessing iteration: (A) Raw, (B) SCT Individual, (C) SCT Whole, and (D) Integrated. For each panel, clustering methods (y-axis) are grouped by the reference label they were evaluated against (Approach, Tissue, or SRA/Sample). Performance is quantified using five clustering metrics, indicated by color: Adjusted Rand Index (ARI), Normalized Mutual Information (NMI), Jaccard Index, Purity, and Variation of Information (VI). Each “Observed” metric value is a single global score calculated by comparing the full set of predicted cluster labels against the true reference labels for all cells. The x-axis shows the Normalized Score, which represents the improvement of a method’s performance over a random baseline. This score is calculated by taking the observed metric value, subtracting the mean value from a set of random clustering permutations, and then scaling the result. For metrics where a higher value is better (ARI, NMI, Jaccard, Purity), the formula is (Observed - Random Mean) / (1 - Random Mean). For Variation of Information (VI), where lower is better, the formula is (Random Mean - Observed) / Random Mean). A score of 1 indicates the best possible performance relative to the random baseline, a score of 0 indicates performance equivalent to random, and negative scores indicate performance worse than random.

A trend emerged when comparing topological and conventional clustering methods across preprocessing iterations. On Raw, SCT Individual, and SCT Whole data, conventional methods like K-Means held a significant performance advantage, with the SCT Whole yielding the highest K-Means performance of all non-integrated methods (ARI_Norm ≈ 0.531). However, this trend inverted sharply after integration. In the Integrated data, hierarchical clustering on landscape distances achieved an ARI_Norm of ≈0.1887, significantly surpassing both Seurat clustering (≈0.0700) and K-Means (≈0.0115). Conventional methods, which rely on local density, may be less effective after integration warps the data’s local geometry. In contrast, PH landscapes capture the global structures that are clarified and enhanced by integration, demonstrating that the integration workflow effectively preprocesses the data to be ideally suited for global topological analysis.

In contrast, other PH-based hierarchical methods, such as those using bottleneck distance or the spectral distance matrix derived from the BDM consistently yielded lower normalized metrics for tissue identification across all iterations. For example, in Raw data, H.Bottleneck_Tissue (ARI_Norm - 0.0021) and H.SDM_Tissue (ARI_Norm 0.0023) performed substantially worse than H.Landscape_Tissue (ARI_Norm 0.0573). This pattern reinforces that not all PH-derived distance matrices contribute equally to biologically meaningful clustering in this multi-tissue context, with landscape-based distances proving most effective among the PH-based hierarchical approaches for discerning tissue types. UMAP visualization further contextualizes these findings (Supplemental Figure 1). When cells in the integrated UMAP space are colored by tissue type (Supplemental Figure 1A), a spectrum of separation is observed. Pancreatic islets and testis form notably distinct and compact clusters, while prostate and substantia nigra also resolve into clear groupings. In contrast, liver and colon appear more diffuse and exhibit considerable intermingling with each other and with the large, overlapping conglomeration of bone marrow and PBMC cells. More strikingly, when the same UMAP is colored by SRA project accession (Supplemental Figure 1B), a clear residual batch effect is visible across the entire embedding. Cells frequently group more strongly by their SRA project of origin than by tissue type, with distinct SRA-specific subclusters evident even within the broader tissue groupings.

The Seurat integration workflow, as applied in this study, was specifically designed to test PH’s capabilities under conditions where technical variation persists. As detailed in the Methods, this involved using SCT Individual normalized data as input to the integration. While the Seurat integration algorithm aims to align shared biological cell states across samples and SRA batches, our design explicitly sought to retain some technical signatures to assess PH’s ability to “see through” or operate robustly in the presence of these batch effects. The UMAP visualization colored by SRA project accession (Supplemental Figure 1B) clearly demonstrates that, as anticipated by this design, substantial SRA-driven structures persist post-integration. These batch-specific signatures appear to contribute significantly to the overall geometry and, consequently, the topology of the integrated data.

This presents a distinct challenge for PH methods. If batch effects are strong enough to impose their own robust topological features, then PH will faithfully detect these batch-driven topological signatures. The task for PH then becomes one of disentangling batch-induced topological features from true, underlying biological topological features. The “topological invariance” of PH means it is robust to deformations that do not change the fundamental connectivity, but if batch effects change this connectivity or create new dominant features, PH will report the topology of this confounded state.

The relatively modest quantitative clustering scores for tissue identification achieved by PH-based methods on the Integrated data likely reflects this complex interplay. The strong SRA-specific clustering suggests that after integration, the topological distances between cells (as measured by BDM, SDM, or LDM) are influenced by their study of origin, in some cases as much or more than by their tissue type. In such a scenario, PH-based clustering might preferentially group samples by SRA project if SRA-specific topological features are more prominent or persistent than tissue-specific ones. Therefore, the success of PH clustering in identifying true biological groupings in integrated data where batch effects are not fully eliminated hinges on the relative strength, persistence, and topological distinctiveness of the biological signal compared to the residual batch noise and its associated topological artifacts. This highlights a critical consideration when applying PH to integrated scRNA-seq datasets.

The choice of clustering algorithm proved to be as critical as the choice of distance metric. In the Raw data, the local-density-based KMeans_Tissue algorithm (ARI_Norm 0.4264) significantly outperformed Spectral.Landscape_ Tissue (ARI_Norm 0.0881), which operates on global topological features. However, in the Integrated data (Supplemental Table 5), Spectral.Landscape_Tissue (ARI_Norm 0.1579, NMI_Norm 0.2358) demonstrated notably strong performance, surpassing KMeans_Tissue (ARI_Norm 0.0115, NMI_Norm 0.0362).

This shift suggests that the integration process, even while leaving some batch effects, may amplify global biological signals that are more effectively captured by the topological landscape representation, enabling spectral embedding to uncover structures that are otherwise obscured by noise and batch effects in unintegrated data. Conversely, in the noisier Raw data, K-means may benefit from its sensitivity to local variance structures. This improvement for tissue identification with Spectral.Landscape post-integration was notable. It is important to acknowledge that this comparison was performed on an integrated dataset generated from inputs where cell-quality covariates (like mitochondrial content) were not regressed out, providing a more challenging scenario for the standard Seurat integration pipeline. Regressing these technical variables prior to integration may improve the performance of conventional clustering methods that are sensitive to such noise. Nonetheless, the strong performance of Spectral.Landscape on this complex dataset is revealing. Its ability to identify not only the tissue-level biological signal but also the residual SRA-level technical signal (ARI_Norm ≈0.1150, NMI_Norm ≈0.3024), significantly outperforming KMeans_SRA (ARI_Norm ≈0.0275, NMI_Norm ≈0.0983) for this task as well (Supplemental Table 5), suggests its general utility in identifying the most prominent global geometric features in transformed data, regardless of their origin.

This inversion of relative performance highlights the complementary roles of the chosen distance representation and the clustering algorithm. It also underscores the potential benefit of combining PH-based representations with spectral clustering, especially for integrated, multi-tissue single-cell datasets. This synergy may offer a promising avenue for improving clustering fidelity and biological relevance, particularly in contexts where subtle, topologically encoded patterns emerge more clearly after integration has mitigated dominant technical effects.

### PH: Betti, Euler, and landscape curve comparisons (bone marrow validation dataset)

A baseline characterization of PH summary behaviors was established on homogeneous data and their responses to various preprocessing strategies with an initial analysis focused on the Betti (β_0_ and β_1_) and Euler characteristic (χ=β_0_ −β_1_) curves derived from the GSE120221 bone marrow dataset collection.

#### Within-iteration analysis (Betti and Euler curves)

Within each data iteration, initial inspection of the individual topological curves (Figure 4) suggested consistent profiles across bone marrow samples, reflecting the tissue’s overall homogeneity. However, a more detailed quantitative analysis of pairwise sample comparisons (Supplemental Table 6) revealed subtle differences and distinct effects from each preprocessing strategy. Across all iterations, KS tests found no significant differences in curve shapes (FDR = 1.0), necessitating the use of Wasserstein distance to capture quantitative variations in inter-sample topology.

**Figure 4:**
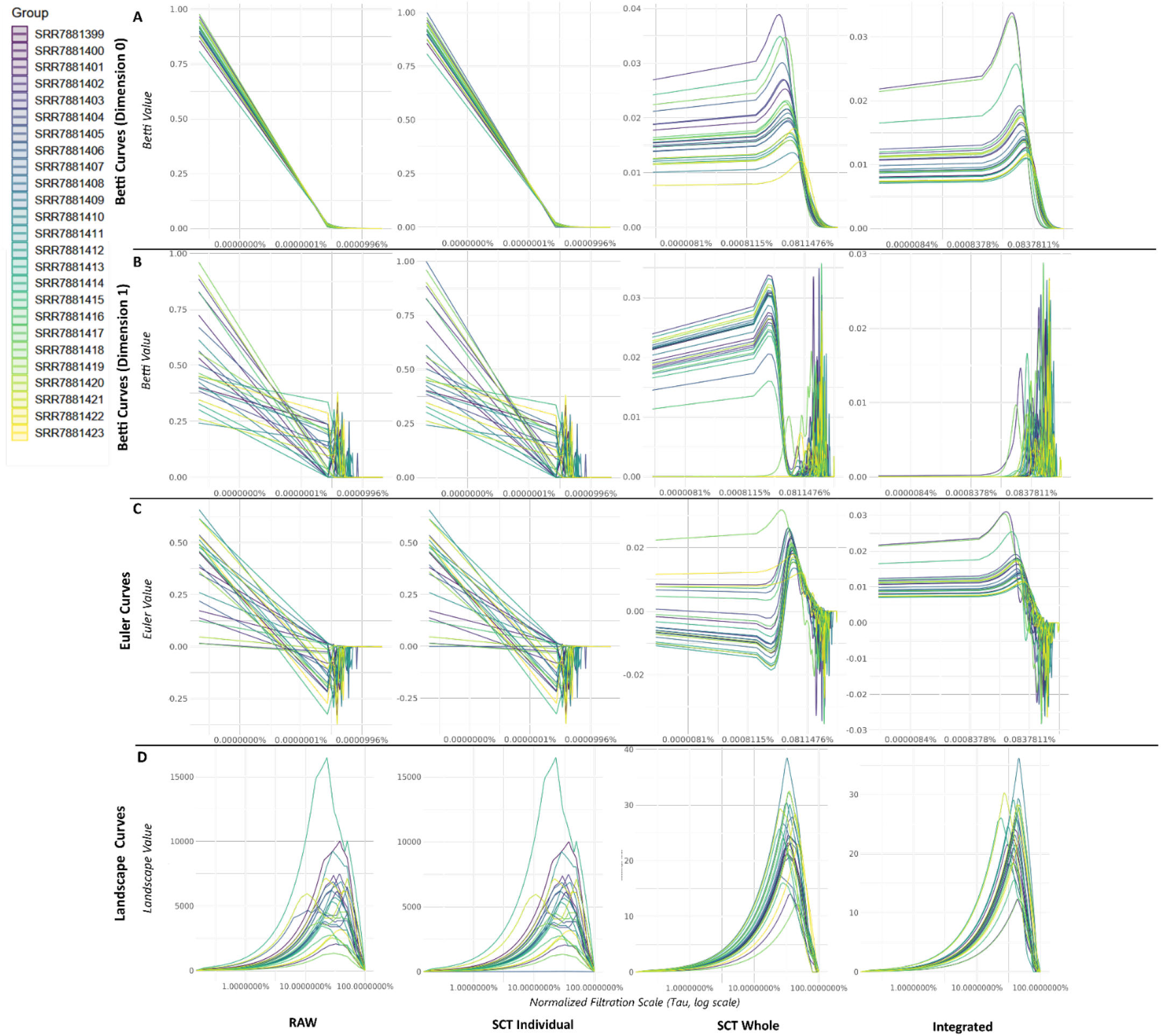
Individual Betti, Euler, and Landscape Curves for Bone Marrow Samples (GSE120221) Across Preprocessing Iterations. The figure displays topological summary curves for each of the 25 individual bone marrow samples from the GSE120221 validation dataset. Each column represents a different data preprocessing iteration: Raw, SCT Individual, SCT Whole, and Integrated. The rows show different topological summaries: (A) Betti 0 (β_0_), (B) Betti 1 (β_1_), (C) Euler characteristic (χ=β_0_−β_1_), and (D) sample level aggregated Landscape curves. The x-axis represents the normalized filtration scale, and the y-axis represents the value of the respective summary. Different colors denote individual samples, identified by their SRA accession numbers as shown in the legend. These plots provide a comparative visualization of the variability in topological signatures among individual samples and how these signatures are visually affected by different normalization and integration strategies.

##### Raw Data

In the Raw data iteration (Figure 4, Column 1), the individual samples exhibited low inter-sample variability. The maximum pairwise Wasserstein distances for β_0_, β_1_, and Euler curves were all modest (≈0.036, ≈0.012, and ≈0.036, respectively). The aggregated landscape curves showed the most variability in this state, with a maximum Wasserstein distance of ≈0.149.

##### SCT Individual

The SCT Individual workflow (Figure 4, Column 2) had a variable effect on homogeneity. While it slightly reduced the maximum Wasserstein distance for β_0_ (≈0.035) and β_1_ (≈0.013) curves, it increased the variability for Euler (≈0.035) and for the landscape curves, where the maximum distance increased to ≈0.130. This indicates individual normalization does not uniformly reduce topological variability and can, in fact, increase the dissimilarity between sample landscapes.

##### SCT Whole

The SCT Whole iteration (Figure 4, Column 3) induced the most dramatic topological changes. A sharp, synchronized β_1_ peak appeared in all samples, leading to a substantial increase in the maximum Wasserstein distance for both β_1_ (≈0.211) and Euler (≈0.081) curves. This suggests the merge-then-normalize approach can introduce significant, coordinated topological artifacts across samples. Interestingly, the landscape curves in this iteration were highly homogeneous, with a low maximum Wasserstein distance (≈0.070).

#### Integrated

The final Integrated iteration (Figure 4, Column 4) demonstrated the most powerful homogenizing effect, but not uniformly across all metrics. The maximum Wasserstein distances for β_0_ (≈0.091), β_1_ (≈0.302), and Euler (≈0.091) curves were the highest of any iteration, indicating that while the mean curve is stabilized (Supplemental Figure 2), the quantitative differences between individual sample curves are amplified. In stark contrast, the landscape curves showed a dramatic convergence: the maximum pairwise Wasserstein distance dropped to just ≈0.08, the lowest of any iteration, demonstrating that integration forces the global landscape representations of the samples into a nearly identical profile. A direct comparison of two representative samples before and after this process (Supplemental Figure 3) illustrates this homogenizing effect.

### Cross-iteration analysis (Betti, Euler, and landscape curves)

Comparing the mean topological signatures (Betti, Euler, and Landscape curves) with confidence intervals, (Figure 5) across preprocessing iterations highlighted the impact of each method.

**Figure 5:**
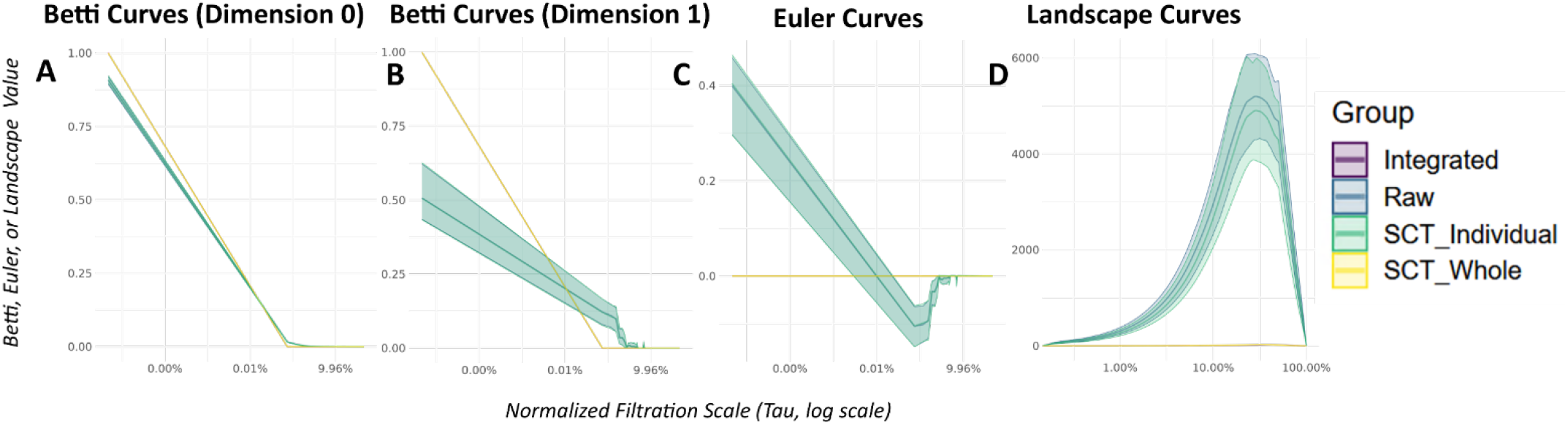
Cross-Iteration Comparison of Mean Topological Summary Curves for the Bone Marrow Validation Set (GSE120221). This figure displays the mean topological summary curves for the entire bone marrow dataset (GSE120221), aggregated to show the overall impact of each preprocessing workflow. Each panel A-D compares the mean signatures across four preprocessing iterations: Raw (blue), SCT Individual (light green), SCT Whole (yellow), and Integrated (purple). Shaded ribbons indicate the 95% confidence interval for each curve. Columns from left to right show: Betti 0 (β_0_) curves, Betti 1 (β_1_) curves, Euler (χ) curves, and aggregated Landscape curves. These plots highlight how the average topological profile of a single homogeneous tissue is morphologically altered by different data processing steps. Quantitative statistical comparisons of the differences between these curves are detailed in Supplemental Table 7. Note: Raw and SCT Individual highly overlap, while the same is true of Integrated and SCT Whole.

The quantitative analysis of mean topological summaries in the bone marrow dataset (Supplemental Table 7) reveals how preprocessing and integration alter the collective topological signature. Direct comparison between the mean Raw and Integrated curves (Figure 5B) showed statistically significant differences for all curve types, with Wasserstein distances of ≈0.0075, ≈0.0034, ≈0.0075, and ≈0.0271, respectively. This demonstrates that the integration workflow, even on this homogeneous dataset, substantially reshapes the average topological profile.

Across all four preprocessing iterations (Figure 5A), the impact of each method was distinct. The mean curves for Raw and SCT Individual were nearly identical, with non-significant KS tests and minimal Wasserstein distances (e.g., ≈0.0023 for landscape curves, p=1.0), suggesting individual normalization had little effect on the average topology. In contrast, both the SCT Whole and Integrated methods induced significant changes relative to the Raw data, particularly in the landscape summary (Wasserstein ≈0.0232 and ≈0.0271, respectively, with p < 0.001 for both).

The contrast between the SCT Whole and SCT Individual normalization strategies is especially pronounced. The mean SCT Whole curves for all summary types were significantly different from the SCT Individual curves (e.g., landscape Wasserstein ≈0.0214, p < 0.001). This divergence, also visible in the sharp, unique peak of the SCT Whole β_1_ curve, suggests the merge-then-normalize approach can introduce distinct topological artifacts not present in individually normalized data.

The subsequent integration step further modified these topological profiles. The mean Integrated landscape curve was significantly different from the SCT Individual curve (Wasserstein ≈0.0266, p < 10^−12^). However, a more complex relationship was observed between the SCT Whole and Integrated states. While the KS test indicated a significant difference in the shape of the landscape curve distributions (FDR < 0.001), the permutation test on their overall Wasserstein distance was only marginally non-significant (Wasserstein ≈0.0198, p ≈ 0.079). This suggests that while integration produced the tightest within-iteration confidence intervals (Supplemental Figure 2), the final mean topological structure is statistically similar in magnitude to that created by the SCT Whole method, despite clear differences in curve shape and smoothness. These findings underscore the importance of assessing preprocessing effects, as different choices can lead to complex and sometimes non-intuitive interpretations.

### PH: Betti, Euler, and landscape curve comparisons (heterogeneous dataset)

Our characterization of PH was extended to a more complex and biologically diverse setting by analyzing a primary dataset collection across eight distinct human tissues Further comparisons were made based on individual Sequence Read Archive (SRA) accessions within each tissue, representing different study origins, and by sequencing approach (scRNA-seq vs. snRNA-seq). Consistent with our validation study, this multi-tissue analysis was conducted across the four standard preprocessing iterations.

Figure 6 presents an overview of mean topological signatures for each of the eight tissues across the preprocessing iterations. These plots provide an initial visualization of how baseline topological structures vary across diverse biological contexts and how these average signatures are modulated by preprocessing and integration. A general observation is that distinct topological profiles are evident across the tissues, particularly in the β_1_, Euler, and Landscape curves, and these profiles undergo noticeable changes with different data processing steps.

**Figure 6:**
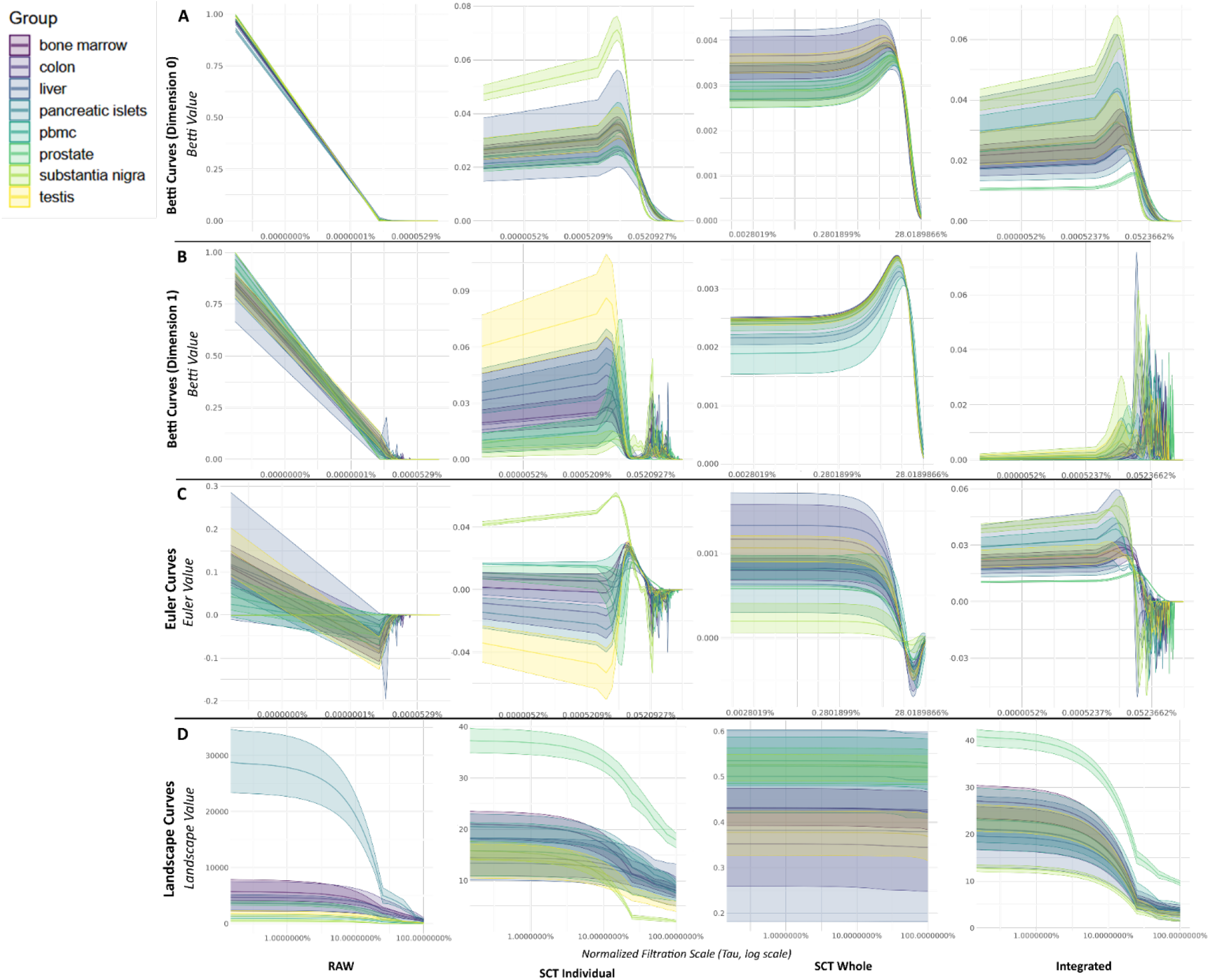
Mean Topological Summary Curves Across Eight Human Tissues and Preprocessing Iterations. The figure displays the mean topological summary curves for each of the eight human tissues in the primary dataset. Each column represents a distinct data preprocessing iteration: Raw, SCT Individual, SCT Whole, and Integrated. The rows show different topological summaries: (A) Betti 0 (β_0_), (B) Betti 1 (β_1_), (C) Euler characteristic (χ=β_0_−β_1_), and (D) aggregated Landscape curves. For each plot, the solid line represents the mean curve calculated across all samples within a given tissue type, and the shaded ribbon indicates the 95% confidence interval derived from bootstrapping. These plots provide a comparative visualization of the average topological signatures across diverse tissues and how these signatures are modulated by different normalization and integration strategies.

#### Within-iteration analysis (Betti, Euler, and landscape curves)

This section details the topological characteristics for each preprocessing iteration, broken down by inter-tissue, inter-study (SRA), and technological comparisons. It draws on mean curve visualizations (Figure 6) and quantitative statistical comparisons (Supplemental Table 8).

### Raw individual iteration

#### Inter-tissue comparisons

In the Raw data iteration (Figure 6, Column 1; Supplemental Table 8), the mean β_0_ curves across the eight tissues generally displayed a monotonic decrease. Pairwise inter-tissue comparisons for these β_0_ curves indicated that while many tissue pairs did not exhibit statistically significant differences in their overall curve shapes after FDR correction (e.g., bone marrow vs. liver, adjusted p≈0.297), several pairs did show significant distinctions (e.g., bone marrow vs. testis, adjusted p≈2.9×10^−9^). The magnitude of these differences, measured by Wasserstein distances, was generally small (e.g., bone marrow vs. colon ≈0.00025). Mean β_1_ curves suggested more pronounced topological variability between tissues. This was substantiated by statistical comparisons, where many of the pairwise tissue comparisons yielded significant adjusted p-values (e.g., bone marrow vs. substantia nigra, adjusted p≈0.003). Wasserstein distances for β_1_ curves reflected these differences (e.g., ≈0.00051 between bone marrow and substantia nigra), quantifying the visual variations in peak structures and overall curve morphologies. Mean Euler curves (χ) also demonstrated significant inter-tissue differences (e.g., bone marrow vs. testis, adjusted p≈2.9×10^−9^, Wasserstein distance ≈0.0011), a distinction apparent in Supplemental Figure 4A. Aggregated landscape curves showed the most pronounced inter-tissue differences in the Raw data. Visually distinct profiles (Figure 6, Row 4, Column 1) were confirmed by statistical testing, which revealed numerous significant pairwise differences (e.g., substantia nigra vs. bone marrow, Wasserstein distance ≈0.185, FDR < 0.001), underscoring their sensitivity to underlying biological structure even before normalization.

#### Inter-study (SRA) comparisons

SRA-level variability (comparisons between different SRA studies) generally showed non-significant shape differences for Betti and Euler curves (most FDR=1.0), though non-zero Wasserstein distances (e.g., β_0_: SRA550660 vs. SRA703206 ≈0.0034) suggested some inter-study variation. Landscape curves at the SRA-level also exhibited variability, with numerous pairwise comparisons showing non-zero Wasserstein distances (e.g., SRA550660 vs. SRA628554, Wass. ≈0.237).

#### Technology comparisons (scRNA-seq vs. snRNA-seq)

Comparisons between scRNA-seq and snRNA-seq approaches revealed statistically significant differences in the shapes of all curve types. For Betti and Euler curves, FDRs were highly significant (e.g., β_0_: FDR≈3.1×10?^6^), as was the difference between landscape curves (FDR≈2.2×10^−7^, Wass. ≈0.148).

### SCT individual iteration

#### Inter-tissue comparisons

Following individual SCTransform normalization (Figure 6, Column 2; Supplementary Table 8), inter-tissue comparisons for mean β_0_ curves continued to show a mix of significant (e.g., bone marrow vs. prostate, FDR≈3.3×10^−12^, Wass. ≈0.032) and non-significant (e.g., bone marrow vs. colon, FDR≈0.091, Wass. ≈0.0021) shape differences. For mean β_1_ curves, significant inter-tissue differences persisted, and Wasserstein distances for some pairs increased compared to Raw data (e.g., bone marrow vs. substantia nigra, FDR≈0.0002, Wass. ≈0.111). The visual separation between tissues like bone marrow and testis remained clear after individual normalization (Supplemental Figure 4B). Mean Euler curves largely mirrored these patterns. For SCT Individual landscape curves, while differences were muted compared to the Raw data, statistical tests still revealed significant distinctions (e.g., bone marrow vs. prostate, FDR≈1.7×10^−12^), though many other pairs became non-significant.

#### Inter-study (SRA) comparisons

SRA-level variability after SCT Individual normalization remained largely non-significant in shape by KS tests, though non-zero Wasserstein distances persisted (e.g., β_0_: SRA550660 vs. SRA703206 ≈0.025). Landscape curves at the SRA-level showed fewer significant differences than in the Raw data, but some non-zero Wasserstein distances remained (max ≈0.00027).

#### Technology comparisons (scRNA-seq vs. snRNA-seq)

The scRNA-seq vs. snRNA-seq comparison continued to show statistically significant differences for β_0_ (FDR≈6.9×10^−5^, Wass. ≈0.031), β_1_ (FDR≈4.7×10^−5^, Wass. ≈0.091), and Euler curves (FDR≈6.9×10^−5^, Wass. ≈0.031). The landscape curve comparison also remained statistically significant (FDR≈5×10^−5^, Wass. ≈0.00016).

### SCT Whole Iteration

#### Inter-tissue comparisons

The SCT Whole iteration (Figure 6, Column 3; Supplementary Table 8), which normalizes the merged dataset, induced topological artifacts. Like the bone marrow validation, this approach created sharp, highly synchronized β_1_ peaks across all eight tissues. This resulted in large Wasserstein distances for β_1_ curves between many tissue pairs (e.g., bone marrow vs. PBMC, Wass. ≈0.28) and numerous significant shape differences (FDR < 0.05). In contrast, the aggregated landscape curves for this iteration were homogeneous, with very low Wasserstein distances between most tissues (e.g., bone marrow vs. colon, Wass. ≈0.0005) and few statistically significant pairwise differences. This highlights a key property of the merge-then-normalize strategy: it can introduce strong, coordinated topological features at the Betti curve level while simultaneously erasing global tissue-specific signatures captured by landscapes (Supplemental Figure 4C).

#### Inter-Study (SRA) and Technology Comparisons

At the SRA level, the SCT Whole iteration showed minimal variability, with most pairwise comparisons being non-significant. However, the technological comparison between scRNA-seq and snRNA-seq revealed highly significant differences (FDR < 10^−5^) with large Wasserstein distances for Betti and Euler curves, indicating this normalization strategy does not resolve technological artifacts.

### Integrated Iteration

#### Inter-tissue comparisons

Data integration using Seurat’s workflow (Figure 6, Column 4; Supplementary Table 8) aimed to harmonize datasets. For mean β_0_ curves, several inter-tissue comparisons remained statistically significant (e.g., bone marrow vs. prostate, FDR≈2.7×10^−12^, Wass. ≈0.046), while others became non-significant, though very close to significant (e.g., bone marrow vs. colon, FDR≈0.051). Mean β_1_ curves often appeared smoother, with many inter-tissue differences preserved or sharpened (e.g., bone marrow vs. testis, FDR≈1.1×10^−5^, Wass. ≈0.024). Mean Euler curves similarly retained significant inter-tissue distinctions (e.g., bone marrow vs. testis, FDR≈1.1×10^−5^, Wass. ≈0.024) (Supplemental Figure 4D). Integrated landscape curves showed that most pairwise inter-tissue landscape comparisons did not have statistically significant shape differences (most FDR>0.05, e.g., bone marrow vs. colon, FDR≈0.386), though some distinctions persisted (e.g., bone marrow vs. prostate, FDR≈0.0017; Wasserstein distances varied, e.g., bone marrow vs. colon ≈0.00018). This suggests that integration does not completely obscure all tissue-specific topological features.

#### Inter-study (SRA) comparisons

SRA-level variability in Integrated data, while largely non-significant by KS tests, still showed non-zero Wasserstein distances (e.g., β_0_: SRA550660 vs. SRA703206 ≈0.030; landscape: SRA550660 vs. SRA645804 ≈0.00027).

#### Technology comparisons (scRNA-seq vs. snRNA-seq)

Comparisons between scRNA-seq and snRNA-seq approaches after integration continued to reveal significant differences for β_0_ (FDR≈6.9×10^−5^, Wass. ≈0.035), β_1_ (FDR≈4.7×10^−5^, Wass. ≈0.123), Euler (FDR≈6.9×10^−5^, Wass. ≈0.035), and landscape curves (FDR≈5×10^−5^, Wass. ≈0.00016).

### Cross-iteration analysis

Directly comparing the mean topological signatures for the entire eight-tissue dataset reveals the global impact of each preprocessing workflow (Figure 7; Supplementary Table 9). In the aggregated view, the mean Raw curves (light green) show a distinct profile. Applying SCT Individual (blue) or SCT Whole (purple) normalization significantly alters these curves, while Integration (pink) produces a final, distinct topological state. The direct comparison between the mean Raw and Integrated curves (Figure 7B) shows statistically significant differences across all summary types (FDR < 10^−15^), with the largest change observed in the landscape summary (Wasserstein ≈ 0.155). This demonstrates that, on a global level, the full preprocessing and integration pipeline fundamentally reshapes the average topological profile of the combined dataset.

**Figure 7:**
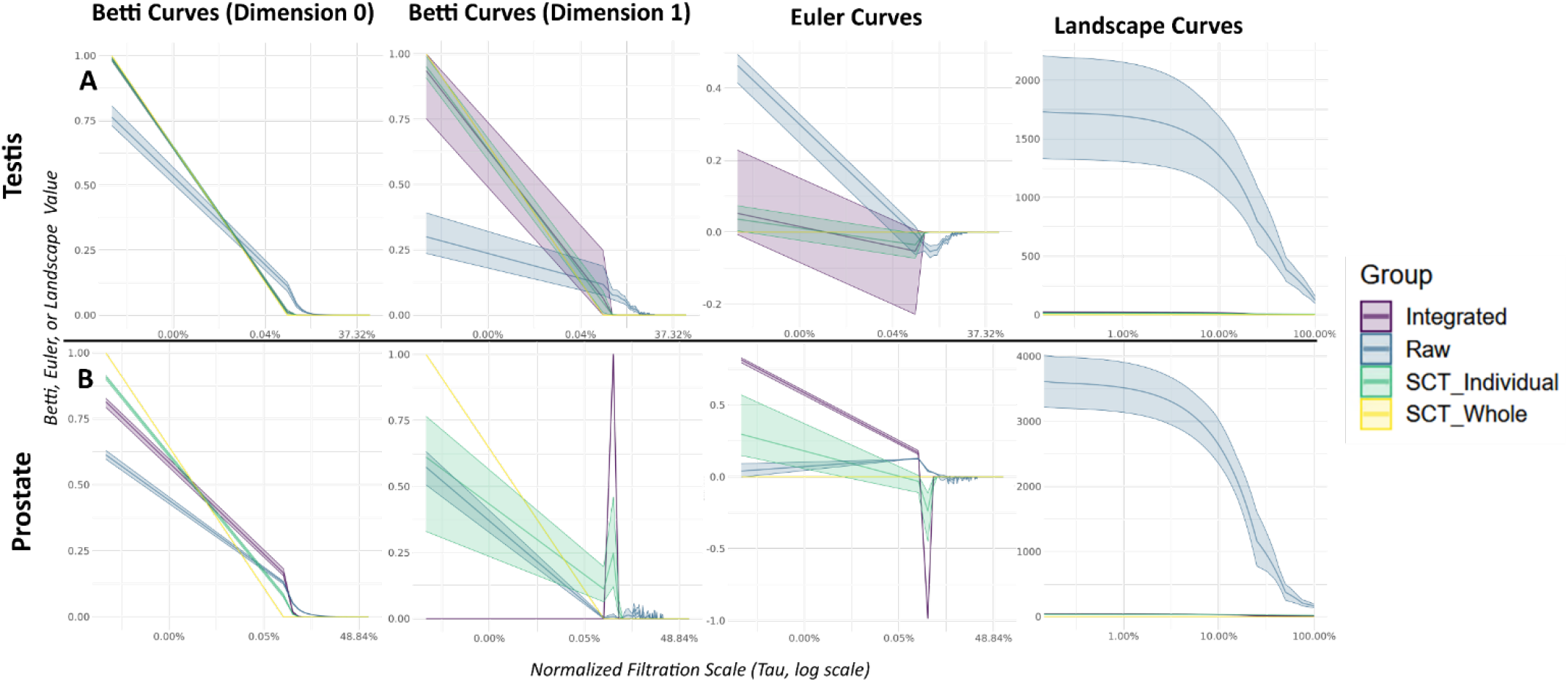
Cross-Iteration Comparison of Mean Topological Summary Curves for Testis and Prostate Tissues. The figure displays the mean topological summary curves for (A) Testis and Prostate tissues, aggregated across all their respective samples to show the overall impact of each preprocessing workflow. Each panel compares the mean signatures across four preprocessing iterations: Raw (blue), SCT Individual (light green), SCT Whole (yellow), and Integrated (purple). Columns from left to right show: Betti 0 (β_0_) curves, Betti 1 (β_1_) curves, Euler (χ) curves, and aggregated Landscape curves. These plots illustrate how the average topological profiles of two distinct tissue types are morphologically altered by different data processing steps. Statistical comparisons of the differences between these curves are detailed in Supplemental Table 9.

The specific impact of these workflows on inter-tissue identification is illustrated in the comparison between bone marrow and testis (Supplemental Figure 4). While the Raw, SCT Individual, and Integrated iterations all maintain a statistically significant topological distinction between the two tissues, the SCT Whole workflow almost completely erases their unique signatures. It forces both tissues into a nearly identical profile characterized by a sharp β1 peak, visually masking their biological differences (Supplemental Figure 4C). This illustrates an example of how a ‘merge-then-normalize’ strategy can introduce technical artifacts that actively obscure true biological heterogeneity.

The impact of preprocessing on intra-tissue signatures is illustrated using testis tissue as an example. The mean β_0_ curve for testis maintains a consistent decay pattern, but confidence intervals narrow from Raw to SCT Individual to Integrated, indicating increased consistency. The mean β_1_ curve shows a broad peak in Raw data, which becomes sharper and slightly higher after SCT Individual normalization (Raw vs. SCT Ind. Wass. ≈ 0.0073, FDR < 10^−13^). Integration further smooths this peak, making it broader again but with a very tight confidence interval, representing a significant change from the SCT Individual state (Wass. ≈ 0.00027, FDR < 0.05). The mean Euler curve reflects these changes, with the initial deep negative deflection becoming smoother and more regular in the Integrated data, accompanied by progressively tighter confidence intervals. The mean landscape curve for testis maintains its overall shape across iterations but undergoes significant quantitative shifts; the transition from Raw to Integrated yields a Wasserstein distance of ≈0.0346 (FDR < 10^−15^), demonstrating that integration reshapes the average landscape profile while increasing its stability.

Analysis of the different sequencing approaches reveals that data processing does not fully harmonize technology-specific topological features. While SCT and Integration workflows alter the curve shapes, significant differences between scRNA-seq and snRNA-seq persist across all iterations. After integration, for example, the mean Euler curves remain significantly different (Wass. ≈ 0.035, FDR≈6.9×10^−5^), as do the landscape curves (Wass. ≈ 0.00016, FDR≈5×10^−5^). This indicates that while integration helps align datasets, fundamental topological differences originating from the sequencing technology are retained.

Broader observations of cross-iteration comparisons for all eight tissues reveal several trends. The transition from Raw to SCT Individual introduces statistically significant changes to the mean curves for most tissues, although the effect size is often modest (e.g., Bone Marrow landscape Wass. ≈0.105, FDR < 10^−15^). In contrast, the comparison between SCT Individual and Integrated data reveals a more complex picture. For tissues like bone marrow and pancreatic islets, the mean Betti and Euler curves show minimal, often non-significant changes, suggesting that for hematopoietic tissues, integration primarily refines the existing topology. However, the landscape curves for these same tissues show significant changes (e.g., Bone Marrow Wass. ≈0.102, FDR < 10?6), indicating that integration is performing a more substantial geometric transformation than is apparent from the Betti curves alone. This highlights the unique sensitivity of landscape summaries to the structural changes induced by data integration.

### Insight for heterogeneous data

Our analysis of the topological characteristics of eight diverse human tissues using PH confirms that distinct tissue-specific topological signatures are discernible, particularly through Betti 1, Euler, and landscape curves (Figure 6). These signatures, however, are variably influenced by common scRNA-seq preprocessing and integration strategies. Raw data provided a baseline where inter-tissue differences were often statistically significant, though SRA-level (inter-study) variations were also present. Individual SCTransform normalization subtly altered these profiles, while the SCT Whole method introduced dramatic, synchronized artifacts across all tissues. In contrast, the Seurat integration workflow generally produced smoother mean topological curves with tighter confidence intervals, indicating an enhancement of within-tissue homogeneity and stability.

The impact of preprocessing on between-tissue distinctions was complex. As illustrated by the comparison between bone marrow and testis (Supplemental Figure 4), the SCT Whole workflow obscured true biological heterogeneity by forcing both tissues into a nearly identical topological profile. Conversely, the SCT Individual and Integrated workflows preserved or even clarified the unique signatures of these distinct tissues. This highlights a critical finding: the normalization strategy can either reveal or conceal biological truth. Furthermore, the persistence of significant topological differences between scRNA-seq and snRNA-seq data even after integration (Supplementary Table 8) underscores PH’s ability to capture structural variations related to RNA source. Clustering performance on this heterogeneous dataset (Figure 3) provided further insights. Conventional methods (Seurat Louvain, K-means) generally outperformed PH-based hierarchical clustering for tissue identification in Raw and SCT Individual data. This indicates that global topological features, when used with hierarchical clustering, may be less effective for broad tissue classification than methods leveraging high-dimensional gene expression variance. However, spectral clustering on landscape-derived distances showed improved performance on Integrated data, surpassing K-means for tissue identification and performing well for resolving SRA identities. This implies that the integration process, while not perfectly removing all SRA-level (batch) effects, might refine the data structure in such a way that landscape-based topological features become more informative for certain clustering algorithms.

The persistence of SRA-level (batch) effects, quantitatively supported by non-zero Wasserstein distances between SRA groups post-integration, is a critical observation. PH, by its nature, is sensitive to the underlying geometry and connectivity of the data. While PH is theoretically invariant to deformations that do not fundamentally alter topology (e.g., stretching or bending), batch effects can introduce more substantial structural changes. If these batch-induced topological features are prominent, PH will detect them. This challenges the notion that PH might serve as a simple panacea for batch effects; rather, its effectiveness is intrinsically tied to the quality of the input data. The Seurat integration method, while effective for many standard analyses, may not optimally prepare the data for PH by fully mitigating topological distortions caused by batch effects while preserving those that are biologically meaningful. The success of PH-based methods in the presence of batch effects therefore hinges on the ability of integration strategies to mitigate these technical distortions in a way that preserves or enhances the biologically relevant topology.

This study demonstrates PH as a valuable framework for characterizing topological diversity in complex biological systems. However, the choice of preprocessing strategy significantly influences the resulting topological summaries. Data integration generally enhances within-group stability, but its impact on between-group distinctions and the mitigation of batch-effect-induced topology requires careful evaluation. These findings highlight the importance of a multi-faceted analytical approach, combining visual inspection of topological curves, quantitative statistical comparisons, and clustering performance, to fully understand the interplay between data processing, batch effects, and the biological signals captured by PH in scRNA-seq studies.

## DISCUSSION

Our analysis reveals the choice of data processing strategy profoundly impacts topological summaries. The central finding is a striking performance inversion: while conventional clustering methods like K-Means are more effective on raw or minimally-processed data, PH-based methods, specifically those using persistence landscapes, significantly outperform them in identifying tissue types after data integration (Figure 3). This suggests a powerful synergy where standard integration algorithms, while potentially disrupting local data structures, effectively clarify the large-scale global topology that landscape-based metrics are uniquely suited to detect. These results position PH not as a replacement for conventional tools, but as a complementary approach for analyzing the global structure of complex, integrated datasets.

A key finding of this study is the inversion of clustering performance between conventional and topological methods following data integration. This phenomenon can be understood by considering how each class of method perceives the data’s geometry, and how integration fundamentally alters that geometry. Conventional algorithms, such as K-Means and Seurat’s graph-based clustering, excel at identifying groups based on local density. They operate by finding dense knots of cells in the high-dimensional space, effectively partitioning the data based on nearest-neighbor relationships. In the Raw and SCT Individual datasets, where biological differences create distinct and relatively compact clusters for each tissue, these methods perform well. However, the Seurat integration workflow, which finds anchors of similar cells across different batches and applies correction vectors to merge them, fundamentally warps the local geometric structure. While this process successfully aligns major cell types on a global scale, it can disrupt the fine-grained neighborhood relationships and local density patterns these conventional methods depend on. This distortion of the local data structure is a likely explanation for the marked decrease in their clustering accuracy post-integration.

In contrast, this geometric transformation appears to create an ideal framework for PH-based analysis. By consolidating the scattered, batch-specific point clouds of each tissue into more coherent “super-clusters,” integration enhances the global separation between biological groups. Persistence landscapes, which quantify the “shape” of data in a way that is less sensitive to local density variations, excel at capturing these large-scale topological features. Instead of focusing on local neighborhoods, landscapes measure the prominence and longevity of overarching structures, such as the distinct connected components formed by each tissue. This creates a synergy whereby the integration workflow removes the topological noise introduced by batch effects, thereby clarifying the global biological structures that landscape-based distance metrics can then effectively distinguish.

These results challenge the notion of PH as a simple panacea for batch effects, instead highlighting its role as a highly sensitive diagnostic tool. The performance of spectral landscape clustering in resolving not only tissue type but also SRA project accession demonstrates that PH faithfully reports the dominant topological features in the data, whether biological or technical. This provides a robust, quantitative method for assessing the quality of integration. While a UMAP may offer insights into residual batch effects, the ability of a PH-based method to cluster by batch provides a clear, data-driven metric of remaining technical variation.

Furthermore, the comparison between scRNA-seq and snRNA-seq data underscores the sensitivity of topological summaries. The Seurat integration workflow, while reducing batch effects, fails to completely harmonize structural differences arising from the sequencing technology. Statistically significant differences between the mean curves for scRNA-seq and snRNA-seq data persist across all topological summaries after integration, including Betti, Euler, and landscape curves (Supplementary Table 8). This indicates that fundamental differences in data structure, likely related to factors like RNA sparsity in snRNA-seq versus scRNA-seq, are retained post-integration. The ability of all PH metrics to detect these residual technological differences highlights their sensitivity, while the distinct patterns captured by landscapes versus Betti/Euler curves showcase their unique perspectives on the data’s geometry.

This study’s findings lead us to refine our initial hypothesis, moving beyond a simple trade-off to a more nuanced understanding of how integration transforms the data’s topological structure. Our results indicate that for local, density-based methods, this transformation can be detrimental; by warping the data to align batches, integration can flatten the fine-grained neighborhood structures that these algorithms depend on, leading to a loss of performance. Conversely, for global topological methods, this transformation is highly beneficial. Integration clarifies the large-scale biological structures that were previously obscured by batch-related noise, creating an ideal substrate for PH landscapes to detect. This reframes the niche for PH in scRNA-seq analysis. It is not a replacement for conventional clustering on raw data, but a powerful tool for analyzing the global structure of complex datasets after they have been harmonized by an integration algorithm.

Comparing a wider range of integration algorithms, such as Harmony or LIGER, could reveal how different geometric transformations of the data impact topological summaries. This could lead to development of topology-aware integration methods explicitly designed to preserve biologically meaningful shape while removing technical artifacts. Applying the validated Integration and landscape workflow to other complex biological problems where global structural changes are paramount represents a promising direction. For instance, in oncology, this approach could characterize intratumor heterogeneity by not only identifying the branching points where new subclones emerge (β_0_ splits) but also by quantifying convergent evolution through the appearance of loop-like features (β_1_). In developmental biology, PH is well-suited to trace continuous cellular trajectories, robustly identifying lineage paths and pinpointing bifurcation events where cell fates diverge, offering a parameter-free method for mapping developmental hierarchies.

## CONCLUSION

We demonstrate that the choice of preprocessing strategy is a non-neutral step in single-cell analysis. It is a fundamental process that reshapes the geometry and topology of the data. Our findings reveal a powerful synergy between Seurat’s anchor-based integration and persistence landscapes, where integration clarifies the global biological structures that landscape-based metrics are uniquely suited to detect. This study establishes PH as a valuable framework for characterizing topological diversity, provides a quantitative tool for assessing the quality of data integration, and underscores the importance of a multi-faceted analytical approach to fully understand the interplay between data processing, batch effects, and the biological signals captured in complex scRNA-seq studies. Ultimately, our results support a refined hypothesis where data integration, rather than masking biological information, transforms the geometric structure of the data. This transformation creates a state where global topological features, best captured by persistence landscapes, are clarified and become more discernible than the local features used by conventional methods.

## ACKNOWLEDGEMENTS

We wish to thank members of the KY INBRE Bioinformatics Core and the SWRM lab members for their helpful insight and feedback.

## DATA AVAILABILITY

All code is available on github at: https://github.com/jcdaneshmand/scPHcompare

## AUTHOR CONTRIBUTIONS

JD: conceptualization, data curation, formal analysis, investigation, methodology, software, validation, visualization, writing – original draft, writing – review & editing. AM: funding acquisition, project administration, resources, supervision, writing – review & editing. JHC: investigation, supervision, writing – review & editing. ECR: conceptualization, funding acquisition, investigation, methodology, project administration, resources, supervision, writing – original draft, writing – review & editing.

