## Supplemental Tables and Figures for "Impact of integration on persistent homology clustering and biological signal detection in scRNA-seq data": Supplementary Materials.docx

**Supplemental Table 1**: Complete scRNA-seq Dataset Collection sourced from PanglaoDB and GEO

| **Sample** | **SRA Number** | **SRS Number** | **Tissue** | **Approach** | **Cells Before Filtering** | **Cells After Filtering** |
| --- | --- | --- | --- | --- | --- | --- |
| SRA621638_SRS2610285 | SRA621638 | SRS2610285 | bone marrow | scRNA-seq | 835 | 832 |
| SRA621638_SRS2610283 | SRA621638 | SRS2610283 | bone marrow | scRNA-seq | 1196 | 1177 |
| SRA740174_SRS3548780 | SRA740174 | SRS3548780 | bone marrow | scRNA-seq | 665 | 614 |
| SRA740174_SRS3548782 | SRA740174 | SRS3548782 | bone marrow | scRNA-seq | 681 | 646 |
| SRA740174_SRS3548783 | SRA740174 | SRS3548783 | bone marrow | scRNA-seq | 1035 | 955 |
| SRA740174_SRS3548781 | SRA740174 | SRS3548781 | bone marrow | scRNA-seq | 1059 | 952 |
| SRA779509_SRS3805264 | SRA779509 | SRS3805264 | bone marrow | scRNA-seq | 992 | 979 |
| SRA779509_SRS3805249 | SRA779509 | SRS3805249 | bone marrow | snRNA-seq | 1133 | 874 |
| SRA779509_SRS3805261 | SRA779509 | SRS3805261 | bone marrow | scRNA-seq | 1829 | 1137 |
| SRA779509_SRS3805267 | SRA779509 | SRS3805267 | bone marrow | scRNA-seq | 3966 | 3427 |
| SRA779509_SRS3805260 | SRA779509 | SRS3805260 | bone marrow | scRNA-seq | 3987 | 2989 |
| SRA779509_SRS3805256 | SRA779509 | SRS3805256 | bone marrow | scRNA-seq | 4958 | 4564 |
| SRA779509_SRS3805253 | SRA779509 | SRS3805253 | bone marrow | scRNA-seq | 5203 | 4111 |
| SRA779509_SRS3805250 | SRA779509 | SRS3805250 | bone marrow | scRNA-seq | 5208 | 3018 |
| SRA779509_SRS3805251 | SRA779509 | SRS3805251 | bone marrow | scRNA-seq | 5345 | 2111 |
| SRA779509_SRS3805257 | SRA779509 | SRS3805257 | bone marrow | scRNA-seq | 5496 | 3831 |
| SRA779509_SRS3805254 | SRA779509 | SRS3805254 | bone marrow | scRNA-seq | 5500 | 4499 |
| SRA779509_SRS3805252 | SRA779509 | SRS3805252 | bone marrow | scRNA-seq | 5540 | 5071 |
| SRA779509_SRS3805268 | SRA779509 | SRS3805268 | bone marrow | scRNA-seq | 5603 | 5042 |
| SRA779509_SRS3805258 | SRA779509 | SRS3805258 | bone marrow | scRNA-seq | 5683 | 5403 |
| SRA779509_SRS3805265 | SRA779509 | SRS3805265 | bone marrow | scRNA-seq | 5735 | 5646 |
| SRA779509_SRS3805269 | SRA779509 | SRS3805269 | bone marrow | scRNA-seq | 5934 | 5226 |
| SRA779509_SRS3805246 | SRA779509 | SRS3805246 | bone marrow | scRNA-seq | 6024 | 5413 |
| SRA779509_SRS3805259 | SRA779509 | SRS3805259 | bone marrow | scRNA-seq | 6032 | 5868 |
| SRA779509_SRS3805263 | SRA779509 | SRS3805263 | bone marrow | scRNA-seq | 6052 | 5200 |
| SRA779509_SRS3805266 | SRA779509 | SRS3805266 | bone marrow | scRNA-seq | 6210 | 5590 |
| SRA779509_SRS3805245 | SRA779509 | SRS3805245 | bone marrow | scRNA-seq | 6248 | 5651 |
| SRA779509_SRS3805248 | SRA779509 | SRS3805248 | bone marrow | scRNA-seq | 6395 | 4536 |
| SRA779509_SRS3805262 | SRA779509 | SRS3805262 | bone marrow | scRNA-seq | 6431 | 6124 |
| SRA779509_SRS3805255 | SRA779509 | SRS3805255 | bone marrow | scRNA-seq | 7559 | 7261 |
| SRA779509_SRS3805247 | SRA779509 | SRS3805247 | bone marrow | scRNA-seq | 7623 | 5416 |
| SRA703206_SRS3296613 | SRA703206 | SRS3296613 | colon | scRNA-seq | 3497 | 2950 |
| SRA703206_SRS3296614 | SRA703206 | SRS3296614 | colon | scRNA-seq | 4651 | 4223 |
| SRA703206_SRS3296611 | SRA703206 | SRS3296611 | colon | scRNA-seq | 4826 | 4175 |
| SRA703206_SRS3296612 | SRA703206 | SRS3296612 | colon | scRNA-seq | 6476 | 5705 |
| SRA728025_SRS3454421 | SRA728025 | SRS3454421 | colon | scRNA-seq | 1883 | 586 |
| SRA728025_SRS3454424 | SRA728025 | SRS3454424 | colon | scRNA-seq | 1898 | 1228 |
| SRA728025_SRS3454423 | SRA728025 | SRS3454423 | colon | scRNA-seq | 2283 | 421 |
| SRA728025_SRS3454422 | SRA728025 | SRS3454422 | colon | scRNA-seq | 2322 | 484 |
| SRA728025_SRS3454426 | SRA728025 | SRS3454426 | colon | scRNA-seq | 3423 | 2009 |
| SRA728025_SRS3454425 | SRA728025 | SRS3454425 | colon | scRNA-seq | 3429 | 1710 |
| SRA728025_SRS3454427 | SRA728025 | SRS3454427 | colon | scRNA-seq | 3840 | 1280 |
| SRA728025_SRS3454430 | SRA728025 | SRS3454430 | colon | scRNA-seq | 4910 | 2127 |
| SRA728025_SRS3454428 | SRA728025 | SRS3454428 | colon | scRNA-seq | 5459 | 1473 |
| SRA716608_SRS3391631 | SRA716608 | SRS3391631 | liver | scRNA-seq | 1425 | 847 |
| SRA716608_SRS3391629 | SRA716608 | SRS3391629 | liver | scRNA-seq | 4190 | 1043 |
| SRA716608_SRS3391630 | SRA716608 | SRS3391630 | liver | scRNA-seq | 4719 | 2284 |
| SRA716608_SRS3391632 | SRA716608 | SRS3391632 | liver | scRNA-seq | 6158 | 554 |
| SRA716608_SRS3391633 | SRA716608 | SRS3391633 | liver | scRNA-seq | 6806 | 1721 |
| SRA760933_SRS3693061 | SRA760933 | SRS3693061 | liver | scRNA-seq | 8030 | 7942 |
| SRA701877_SRS3279688 | SRA701877 | SRS3279688 | pancreatic islets | snRNA-seq | 442 | 396 |
| SRA701877_SRS3279685 | SRA701877 | SRS3279685 | pancreatic islets | scRNA-seq | 1123 | 943 |
| SRA701877_SRS3279689 | SRA701877 | SRS3279689 | pancreatic islets | scRNA-seq | 1245 | 1090 |
| SRA701877_SRS3279691 | SRA701877 | SRS3279691 | pancreatic islets | scRNA-seq | 1794 | 1640 |
| SRA701877_SRS3279686 | SRA701877 | SRS3279686 | pancreatic islets | scRNA-seq | 1805 | 1632 |
| SRA701877_SRS3279684 | SRA701877 | SRS3279684 | pancreatic islets | scRNA-seq | 2052 | 1885 |
| SRA701877_SRS3279687 | SRA701877 | SRS3279687 | pancreatic islets | scRNA-seq | 2416 | 2106 |
| SRA701877_SRS3279690 | SRA701877 | SRS3279690 | pancreatic islets | scRNA-seq | 2492 | 2157 |
| SRA701877_SRS3279694 | SRA701877 | SRS3279694 | pancreatic islets | scRNA-seq | 2499 | 2213 |
| SRA701877_SRS3279692 | SRA701877 | SRS3279692 | pancreatic islets | scRNA-seq | 2884 | 2619 |
| SRA701877_SRS3279693 | SRA701877 | SRS3279693 | pancreatic islets | snRNA-seq | 2953 | 2491 |
| SRA701877_SRS3279695 | SRA701877 | SRS3279695 | pancreatic islets | snRNA-seq | 3508 | 2992 |
| SRA550660_SRS2089636 | SRA550660 | SRS2089636 | pbmc | scRNA-seq | 1580 | 930 |
| SRA550660_SRS2089638 | SRA550660 | SRS2089638 | pbmc | scRNA-seq | 1818 | 1210 |
| SRA550660_SRS2089635 | SRA550660 | SRS2089635 | pbmc | scRNA-seq | 1860 | 1138 |
| SRA550660_SRS2089637 | SRA550660 | SRS2089637 | pbmc | snRNA-seq | 10292 | 9071 |
| SRA550660_SRS2089639 | SRA550660 | SRS2089639 | pbmc | snRNA-seq | 10940 | 8839 |
| SRA628554_SRS2664365 | SRA628554 | SRS2664365 | pbmc | snRNA-seq | 3081 | 3141 |
| SRA628554_SRS2664364 | SRA628554 | SRS2664364 | pbmc | snRNA-seq | 3352 | 3410 |
| SRA713577_SRS3363004 | SRA713577 | SRS3363004 | pbmc | scRNA-seq | 3334 | 3326 |
| SRA749327_SRS3693912 | SRA749327 | SRS3693912 | pbmc | scRNA-seq | 1352 | 945 |
| SRA749327_SRS3693911 | SRA749327 | SRS3693911 | pbmc | scRNA-seq | 1536 | 935 |
| SRA749327_SRS3693910 | SRA749327 | SRS3693910 | pbmc | scRNA-seq | 1929 | 1889 |
| SRA749327_SRS3693913 | SRA749327 | SRS3693913 | pbmc | scRNA-seq | 3744 | 2644 |
| SRA742961_SRS3565205 | SRA742961 | SRS3565205 | prostate | scRNA-seq | 4813 | 4751 |
| SRA742961_SRS3565204 | SRA742961 | SRS3565204 | prostate | scRNA-seq | 5198 | 4900 |
| **Sample** | **SRA Number** | **SRS Number** | **Tissue** | **Approach** | **Cells Before Filtering** | **Cells After Filtering** |
| SRA742961_SRS3565202 | SRA742961 | SRS3565202 | prostate | scRNA-seq | 5804 | 5748 |
| SRA742961_SRS3565210 | SRA742961 | SRS3565210 | prostate | scRNA-seq | 5923 | 5566 |
| SRA742961_SRS3565195 | SRA742961 | SRS3565195 | prostate | scRNA-seq | 6101 | 5718 |
| SRA742961_SRS3565211 | SRA742961 | SRS3565211 | prostate | scRNA-seq | 6264 | 5981 |
| SRA742961_SRS3565207 | SRA742961 | SRS3565207 | prostate | scRNA-seq | 6323 | 6257 |
| SRA742961_SRS3565203 | SRA742961 | SRS3565203 | prostate | scRNA-seq | 6453 | 6335 |
| SRA742961_SRS3565206 | SRA742961 | SRS3565206 | prostate | scRNA-seq | 6950 | 6956 |
| SRA742961_SRS3565209 | SRA742961 | SRS3565209 | prostate | scRNA-seq | 7132 | 7044 |
| SRA742961_SRS3565201 | SRA742961 | SRS3565201 | prostate | scRNA-seq | 7479 | 7482 |
| SRA742961_SRS3565198 | SRA742961 | SRS3565198 | prostate | scRNA-seq | 7718 | 7396 |
| SRA742961_SRS3565196 | SRA742961 | SRS3565196 | prostate | scRNA-seq | 7789 | 7791 |
| SRA742961_SRS3565199 | SRA742961 | SRS3565199 | prostate | scRNA-seq | 8069 | 8050 |
| SRA742961_SRS3565208 | SRA742961 | SRS3565208 | prostate | snRNA-seq | 8538 | 8118 |
| SRA742961_SRS3565197 | SRA742961 | SRS3565197 | prostate | scRNA-seq | 11468 | 11475 |
| SRA850958_SRS4386101 | SRA850958 | SRS4386101 | substantia nigra | snRNA-seq | 906 | 1015 |
| SRA850958_SRS4386110 | SRA850958 | SRS4386110 | substantia nigra | scRNA-seq | 1065 | 617 |
| SRA850958_SRS4386109 | SRA850958 | SRS4386109 | substantia nigra | scRNA-seq | 1387 | 809 |
| SRA850958_SRS4386113 | SRA850958 | SRS4386113 | substantia nigra | snRNA-seq | 1973 | 584 |
| SRA850958_SRS4386112 | SRA850958 | SRS4386112 | substantia nigra | snRNA-seq | 2432 | 713 |
| SRA850958_SRS4386111 | SRA850958 | SRS4386111 | substantia nigra | snRNA-seq | 2590 | 778 |
| SRA645804_SRS2823411 | SRA645804 | SRS2823411 | testis | scRNA-seq | 2167 | 1031 |
| SRA645804_SRS2823405 | SRA645804 | SRS2823405 | testis | scRNA-seq | 3598 | 3680 |
| SRA645804_SRS2823406 | SRA645804 | SRS2823406 | testis | scRNA-seq | 3989 | 4297 |
| SRA645804_SRS2823410 | SRA645804 | SRS2823410 | testis | snRNA-seq | 4045 | 3561 |
| SRA645804_SRS2823407 | SRA645804 | SRS2823407 | testis | scRNA-seq | 4046 | 2708 |
| SRA645804_SRS2823404 | SRA645804 | SRS2823404 | testis | scRNA-seq | 4197 | 4205 |
| SRA645804_SRS2823408 | SRA645804 | SRS2823408 | testis | snRNA-seq | 4306 | 3006 |
| SRA645804_SRS3572594 | SRA645804 | SRS3572594 | testis | snRNA-seq | 4574 | 4035 |
| SRA645804_SRS2823409 | SRA645804 | SRS2823409 | testis | snRNA-seq | 4791 | 3897 |
| SRA645804_SRS2823412 | SRA645804 | SRS2823412 | testis | snRNA-seq | 5299 | 3846 |
| SRA667709_SRS3065426 | SRA667709 | SRS3065426 | testis | scRNA-seq | 2777 | 1118 |
| SRA667709_SRS3065428 | SRA667709 | SRS3065428 | testis | scRNA-seq | 3007 | 1315 |
| SRA667709_SRS3065427 | SRA667709 | SRS3065427 | testis | scRNA-seq | 3045 | 1945 |
| SRA667709_SRS3065429 | SRA667709 | SRS3065429 | testis | scRNA-seq | 3066 | 1847 |
| SRA667709_SRS3065431 | SRA667709 | SRS3065431 | testis | scRNA-seq | 3586 | 1410 |
| SRA667709_SRS3065430 | SRA667709 | SRS3065430 | testis | scRNA-seq | 4020 | 1486 |
| SRA784242_SRS3822686 | SRA784242 | SRS3822686 | testis | scRNA-seq | 1580 | 623 |
| SRA784242_SRS3822683 | SRA784242 | SRS3822683 | testis | scRNA-seq | 1631 | 647 |
| SRA784242_SRS3822682 | SRA784242 | SRS3822682 | testis | scRNA-seq | 2455 | 574 |
| SRA784242_SRS3822680 | SRA784242 | SRS3822680 | testis | scRNA-seq | 2487 | 537 |
| SRA826293_SRS4181124 | SRA826293 | SRS4181124 | testis | scRNA-seq | 3307 | 1664 |
| SRA826293_SRS4181125 | SRA826293 | SRS4181125 | testis | scRNA-seq | 3931 | 609 |
| SRA826293_SRS4181123 | SRA826293 | SRS4181123 | testis | scRNA-seq | 4876 | 4490 |
| SRA826293_SRS4181126 | SRA826293 | SRS4181126 | testis | scRNA-seq | 5464 | 2250 |
| SRA826293_SRS4181127 | SRA826293 | SRS4181127 | testis | scRNA-seq | 5705 | 5276 |
| SRA826293_SRS4181128 | SRA826293 | SRS4181128 | testis | scRNA-seq | 6198 | 4681 |
| SRA826293_SRS4181129 | SRA826293 | SRS4181129 | testis | scRNA-seq | 7860 | 4038 |
| SRA826293_SRS4181130 | SRA826293 | SRS4181130 | testis | scRNA-seq | 9028 | 7965 |

**Supplemental Table 2**: For Validation: Bone Marrow dataset collection (GSE120221)

| **GEO Accession / Sample Name** | **Run** | **Cells Before Filtering** | **Cells After Filtering** |
| --- | --- | --- | --- |
| GSM3396161 | SRR7881399 | 2994 | 2492 |
| GSM3396162 | SRR7881400 | 3293 | 2832 |
| GSM3396163 | SRR7881401 | 3556 | 1831 |
| GSM3396164 | SRR7881402 | 1052 | 789 |
| GSM3396165 | SRR7881403 | 3136 | 1632 |
| GSM3396166 | SRR7881404 | 3939 | 1940 |
| GSM3396167 | SRR7881405 | 3746 | 3361 |
| GSM3396168 | SRR7881406 | 4283 | 1500 |
| GSM3396169 | SRR7881407 | 4516 | 3507 |
| GSM3396170 | SRR7881408 | 3446 | 2663 |
| GSM3396171 | SRR7881409 | 7247 | 6990 |
| GSM3396172 | SRR7881410 | 4548 | 4257 |
| GSM3396173 | SRR7881411 | 3964 | 2483 |
| GSM3396174 | SRR7881412 | 4522 | 4279 |
| GSM3396175 | SRR7881413 | 5013 | 4925 |
| GSM3396176 | SRR7881414 | 3383 | 3323 |
| GSM3396177 | SRR7881415 | 1700 | 1020 |
| GSM3396178 | SRR7881416 | 3593 | 2601 |
| GSM3396179 | SRR7881417 | 2437 | 1740 |
| GSM3396180 | SRR7881418 | 1138 | 986 |
| GSM3396181 | SRR7881419 | 2367 | 1780 |
| GSM3396182 | SRR7881420 | 4726 | 4639 |
| GSM3396183 | SRR7881421 | 4293 | 4050 |
| GSM3396184 | SRR7881422 | 4118 | 3677 |
| GSM3396185 | SRR7881423 | 3643 | 3097 |

**Supplemental Table 3**: R Package Versions

| **Package** | **Version** |
| --- | --- |
| aricode | 1.0.3 |
| BiocSingular | 1.20.0 |
| circlize | 0.4.16 |
| ComplexHeatmap | 2.20.0 |
| dendextend | 1.19.0 |
| digest | 0.6.37 |
| doParallel | 1.0.17 |
| foreach | 1.5.2 |
| gridExtra | 2.3 |
| igraph | 2.1.4 |
| kernlab | 0.9-33 |
| mclust | 6.1.1 |
| pheatmap | 1.0.12 |
| plyr | 1.8.9 |
| RColorBrewer | 1.1-3 |
| reshape2 | 1.4.4 |
| ripserr | 0.3.0 |
| Rtsne | 0.17 |
| scCustomize | 3.0.1 |
| Seurat | 5.3.0 |
| SeuratDisk | 0.0.0.9021 |
| SeuratObject | 5.0.2 |
| stringr | 1.5.1 |
| TDA | 1.9.4 |
| TDAstats | 0.4.1 |
| tidyverse | 2.0.0 |
| transport | 0.15-4 |
| viridis | 0.6.5 |
| Matrix | 1.7-3 |
| patchwork | 1.3.0 |
| parallel | 4.4.1 |

**Supplemental Table 4**: Bone Marrow Normalized Cluster Comparison Results

|  |  |  | **ARI** |  | **NMI** |  | **Jaccard** |  | **VI** |  | **Purity** |  |
| --- | --- | --- | --- | --- | --- | --- | --- | --- | --- | --- | --- | --- |
| **Iteration** | **Method** | **Reference**  **Col** | **Norm** | **FDR** | **Norm** | **FDR** | **Norm** | **FDR** | **Norm** | **FDR** | **Norm** | **FDR** |
| Raw | seurat_raw_sra | orig.ident | 0.0337 | 0 | 0.1116 | 0 | -0.0286 | 1 | -0.2073 | 1 | -0.2582 | 1 |
| Raw | seurat_raw_tissue | Tissue | -0.0431 | NA | -0.0553 | NA | -0.1104 | NA | 1 | NA | -0.5761 | NA |
| Raw | seurat_raw_approach | Approach | -0.0431 | NA | -0.0553 | NA | -0.1104 | NA | 1 | NA | -0.5761 | NA |
| Raw | kmeans_raw_tissue | Tissue | 0 | NA | 0 | NA | -0.2665 | NA | 1 | NA | -0.3561 | NA |
| Raw | kmeans_raw_sra | orig.ident | 0.0323 | 0 | 0.103 | 0 | -0.0141 | 1 | -0.2116 | 1 | -0.2592 | 1 |
| Raw | kmeans_raw_approach | Approach | 0 | NA | 0 | NA | -0.2665 | NA | 1 | NA | -0.3561 | NA |
| Raw | hierarchical_bdm_ph_raw_tissue | Tissue | 0 | NA | 0 | NA | -0.2665 | NA | 1 | NA | -0.3561 | NA |
| Raw | hierarchical_sdm_ph_raw_tissue | Tissue | 0 | NA | 0 | NA | -0.2665 | NA | 1 | NA | -0.3561 | NA |
| Raw | hierarchical_landscape_ph_raw_tissue | Tissue | 0 | NA | 0 | NA | -0.2665 | NA | 1 | NA | -0.3561 | NA |
| Raw | hierarchical_bdm_ph_raw_sra | orig.ident | 1 | 0 | 1 | 0 | 1 | 0 | 1 | 0 | 0 | 1 |
| Raw | hierarchical_sdm_ph_raw_sra | orig.ident | 1 | 0 | 1 | 0 | 1 | 0 | 1 | 0 | 0 | 1 |
| Raw | hierarchical_landscape_ph_raw_sra | orig.ident | 1 | 0 | 1 | 0 | 1 | 0 | 1 | 0 | 0 | 1 |
| Raw | hierarchical_bdm_ph_raw_approach | Approach | 0 | NA | 0 | NA | -0.2665 | NA | 1 | NA | -0.3561 | NA |
| Raw | hierarchical_sdm_ph_raw_approach | Approach | 0 | NA | 0 | NA | -0.2665 | NA | 1 | NA | -0.3561 | NA |
| Raw | hierarchical_landscape_ph_raw_approach | Approach | 0 | NA | 0 | NA | -0.2665 | NA | 1 | NA | -0.3561 | NA |
| SCT Individual | seurat_sct_individual_sra | orig.ident | 0.0861 | 0 | 0.2422 | 0 | 0.0211 | 0.125 | -0.0439 | 1 | -0.294 | 1 |
| SCT Individual | seurat_sct_individual_tissue | Tissue | -0.0809 | NA | -0.1353 | NA | -0.1041 | NA | 1 | NA | -0.8603 | NA |
| SCT Individual | seurat_sct_individual_approach | Approach | -0.0809 | NA | -0.1353 | NA | -0.1041 | NA | 1 | NA | -0.8603 | NA |
| SCT Individual | kmeans_sct_individual_tissue | Tissue | 0 | NA | 0 | NA | -0.2665 | NA | 1 | NA | -0.3561 | NA |
| SCT Individual | kmeans_sct_individual_sra | orig.ident | 0.0382 | 0 | 0.1491 | 0 | -0.0175 | 1 | -0.1545 | 1 | -0.287 | 1 |
| SCT Individual | kmeans_sct_individual_approach | Approach | 0 | NA | 0 | NA | -0.2665 | NA | 1 | NA | -0.3561 | NA |
| SCT Individual | hierarchical_bdm_ph_sct_individual_tissue | Tissue | 0 | NA | 0 | NA | -0.2665 | NA | 1 | NA | -0.3561 | NA |
| SCT Individual | hierarchical_sdm_ph_sct_individual_tissue | Tissue | 0 | NA | 0 | NA | -0.2665 | NA | 1 | NA | -0.3561 | NA |
| SCT Individual | hierarchical_landscape_ph_sct_individual_tissue | Tissue | 0 | NA | 0 | NA | -0.2665 | NA | 1 | NA | -0.3561 | NA |
| SCT Individual | hierarchical_bdm_ph_sct_individual_sra | orig.ident | 1 | 0 | 1 | 0 | 1 | 0 | 1 | 0 | 0 | 1 |
| SCT Individual | hierarchical_sdm_ph_sct_individual_sra | orig.ident | 1 | 0 | 1 | 0 | 1 | 0 | 1 | 0 | 0 | 1 |
| SCT Individual | hierarchical_landscape_ph_sct_individual_sra | orig.ident | 1 | 0 | 1 | 0 | 1 | 0 | 1 | 0 | 0 | 1 |
| SCT Individual | hierarchical_bdm_ph_sct_individual_approach | Approach | 0 | NA | 0 | NA | -0.2665 | NA | 1 | NA | -0.3561 | NA |
| SCT Individual | hierarchical_sdm_ph_sct_individual_approach | Approach | 0 | NA | 0 | NA | -0.2665 | NA | 1 | NA | -0.3561 | NA |
| SCT Individual | hierarchical_landscape_ph_sct_individual_approach | Approach | 0 | NA | 0 | NA | -0.2665 | NA | 1 | NA | -0.3561 | NA |
| SCT Whole | seurat_sct_whole_sra | orig.ident | 0.0562 | 0 | 0.1944 | 0 | 0.0018 | 0.5 | -0.0984 | 1 | -0.2863 | 1 |
| SCT Whole | seurat_sct_whole_tissue | Tissue | -0.0522 | NA | -0.0996 | NA | -0.0906 | NA | 1 | NA | -0.7309 | NA |
| SCT Whole | seurat_sct_whole_approach | Approach | -0.0522 | NA | -0.0996 | NA | -0.0906 | NA | 1 | NA | -0.7309 | NA |
| SCT Whole | kmeans_sct_whole_tissue | Tissue | 0 | NA | 0 | NA | -0.2665 | NA | 1 | NA | -0.3561 | NA |
| SCT Whole | kmeans_sct_whole_sra | orig.ident | 0.0432 | 0 | 0.134 | 0 | -0.001 | 0.8 | -0.1684 | 1 | -0.2769 | 1 |
| SCT Whole | kmeans_sct_whole_approach | Approach | 0 | NA | 0 | NA | -0.2665 | NA | 1 | NA | -0.3561 | NA |
| SCT Whole | hierarchical_bdm_ph_sct_whole_tissue | Tissue | 0 | NA | 0 | NA | -0.2665 | NA | 1 | NA | -0.3561 | NA |
| SCT Whole | hierarchical_sdm_ph_sct_whole_tissue | Tissue | 0 | NA | 0 | NA | -0.2665 | NA | 1 | NA | -0.3561 | NA |
| SCT Whole | hierarchical_landscape_ph_sct_whole_tissue | Tissue | 0 | NA | 0 | NA | -0.2665 | NA | 1 | NA | -0.3561 | NA |
| SCT Whole | hierarchical_bdm_ph_sct_whole_sra | orig.ident | 1 | 0 | 1 | 0 | 1 | 0 | 1 | 0 | 0 | 1 |
| SCT Whole | hierarchical_sdm_ph_sct_whole_sra | orig.ident | 1 | 0 | 1 | 0 | 1 | 0 | 1 | 0 | 0 | 1 |
| SCT Whole | hierarchical_landscape_ph_sct_whole_sra | orig.ident | 1 | 0 | 1 | 0 | 1 | 0 | 1 | 0 | 0 | 1 |
| SCT Whole | hierarchical_bdm_ph_sct_whole_approach | Approach | 0 | NA | 0 | NA | -0.2665 | NA | 1 | NA | -0.3561 | NA |
| SCT Whole | hierarchical_sdm_ph_sct_whole_approach | Approach | 0 | NA | 0 | NA | -0.2664 | NA | 1 | NA | -0.3561 | NA |
| SCT Whole | hierarchical_landscape_ph_sct_whole_approach | Approach | 0 | NA | 0 | NA | -0.2665 | NA | 1 | NA | -0.3561 | NA |
| Integrated | seurat_integrated_sra | orig.ident | 0.0151 | 0 | 0.0588 | 0 | -0.029 | 1 | -0.2726 | 1 | -0.2332 | 1 |
| Integrated | seurat_integrated_tissue | Tissue | -0.0118 | NA | -0.0263 | NA | -0.0792 | NA | 1 | NA | -0.4872 | NA |
| Integrated | seurat_integrated_approach | Approach | -0.0118 | NA | -0.0263 | NA | -0.0792 | NA | 1 | NA | -0.4872 | NA |
| Integrated | kmeans_integrated_tissue | Tissue | 0 | NA | 0 | NA | -0.2665 | NA | 1 | NA | -0.3561 | NA |
| Integrated | kmeans_integrated_sra | orig.ident | 0.01 | 0 | 0.0481 | 0 | -0.0429 | 1 | -0.29 | 1 | -0.2336 | 1 |
| Integrated | kmeans_integrated_approach | Approach | 0 | NA | 0 | NA | -0.2665 | NA | 1 | NA | -0.3561 | NA |
| Integrated | hierarchical_bdm_ph_integrated_tissue | Tissue | 0 | NA | 0 | NA | -0.2665 | NA | 1 | NA | -0.3561 | NA |
|  |  |  | **ARI** |  | **NMI** |  | **Jaccard** |  | **VI** |  | **Purity** |  |
| **Iteration** | **Method** | **Reference**  **Col** | **Norm** | **FDR** | **Norm** | **FDR** | **Norm** | **FDR** | **Norm** | **FDR** | **Norm** | **FDR** |
| Integrated | hierarchical_sdm_ph_integrated_tissue | Tissue | 0 | NA | 0 | NA | -0.2665 | NA | 1 | NA | -0.3561 | NA |
| Integrated | hierarchical_landscape_ph_integrated_tissue | Tissue | 0 | NA | 0 | NA | -0.2665 | NA | 1 | NA | -0.3561 | NA |
| Integrated | hierarchical_bdm_ph_integrated_sra | orig.ident | 1 | 0 | 1 | 0 | 1 | 0 | 1 | 0 | 0 | 1 |
| Integrated | hierarchical_sdm_ph_integrated_sra | orig.ident | 1 | 0 | 1 | 0 | 1 | 0 | 1 | 0 | 0 | 1 |
| Integrated | hierarchical_landscape_ph_integrated_sra | orig.ident | 1 | 0 | 1 | 0 | 1 | 0 | 1 | 0 | 0 | 1 |
| Integrated | hierarchical_bdm_ph_integrated_approach | Approach | 0 | NA | 0 | NA | -0.2665 | NA | 1 | NA | -0.3561 | NA |
| Integrated | hierarchical_sdm_ph_integrated_approach | Approach | 0 | NA | 0 | NA | -0.2665 | NA | 1 | NA | -0.3561 | NA |
| Integrated | hierarchical_landscape_ph_integrated_approach | Approach | 0 | NA | 0 | NA | -0.2665 | NA | 1 | NA | -0.3561 | NA |

**Supplemental Table 5**: Multi-Tissue Dataset Collection Normalized Cluster Comparison Results

|  |  |  | **ARI** |  | **NMI** |  | **Jaccard** |  | **VI** |  | **Purity** |  |
| --- | --- | --- | --- | --- | --- | --- | --- | --- | --- | --- | --- | --- |
| **Iteration** | **Method** | **ReferenceCol** | **Norm** | **FDR** | **Norm** | **FDR** | **Norm** | **FDR** | **Norm** | **FDR** | **Norm** | **FDR** |
| Raw | seurat_raw_sra | SRA | 0.0269 | 0 | 0.2021 | 0 | 0.0143 | 0 | 0.2021 | 0 | 0 | 1 |
| Raw | seurat_raw_tissue | Tissue | -0.0003 | 1 | 0.0525 | 0 | -0.0002 | 0.9333 | 0.0525 | 0 | 0 | 1 |
| Raw | seurat_raw_approach | Approach | -0.1031 | 1 | -0.5218 | 1 | -0.0516 | 1 | -0.5218 | 1 | 0 | 1 |
| Raw | kmeans_raw_tissue | Tissue | 0.4264 | 0 | 0.4576 | 0 | 0.3001 | 0 | 0.4787 | 0 | 0.5347 | 0 |
| Raw | kmeans_raw_sra | SRA | 0.389 | 0 | 0.5605 | 0 | 0.252 | 0 | 0.4891 | 0 | 0.6025 | 0 |
| Raw | kmeans_raw_approach | Approach | -0.0166 | 1 | -0.0025 | 1 | 0.9029 | 0 | 0.8879 | 0 | 0.9853 | 0 |
| Raw | hierarchical_bdm_ph_raw_tissue | Tissue | -0.0021 | 1 | 0.0243 | 0 | -0.001 | 0.8471 | -0.0197 | 1 | 0.0636 | 0 |
| Raw | hierarchical_sdm_ph_raw_tissue | Tissue | 0.0023 | 1 | 0.0012 | 0.6 | 0.002 | 0.32 | -0.0785 | 1 | 0.0773 | 0 |
| Raw | hierarchical_landscape_ph_raw_tissue | Tissue | 0.0573 | 0 | 0.0817 | 0 | 0.0346 | 0 | 0.0516 | 0 | 0.1059 | 0 |
| Raw | hierarchical_bdm_ph_raw_sra | SRA | -0.03 | 1 | 0.0586 | 0 | -0.0380 | 1 | -0.0783 | 1 | 0.0076 | 0.6316 |
| Raw | hierarchical_sdm_ph_raw_sra | SRA | -0.0105 | 1 | 0.0167 | 0.1714 | -0.0397 | 1 | -0.139 | 1 | 0.0982 | 0 |
| Raw | hierarchical_landscape_ph_raw_sra | SRA | 0.1077 | 0 | 0.2515 | 0 | 0.0521 | 0 | 0.111 | 0 | 0.1641 | 0 |
| Raw | hierarchical_bdm_ph_raw_approach | Approach | -0.0128 | 1 | -0.0147 | 1 | 0.931 | 0 | 0.9096 | 0 | 0.9852 | 0 |
| Raw | hierarchical_sdm_ph_raw_approach | Approach | -0.0073 | 1 | -0.0044 | 1 | 0.9597 | 0 | 0.9422 | 0 | 0.9854 | 0 |
| Raw | hierarchical_landscape_ph_raw_approach | Approach | -0.0280 | 1 | -0.0189 | 1 | 0.6755 | 0 | 0.759 | 0 | 0.9848 | 0 |
| Raw | spectral_bdm_raw_tissue | Tissue | -0.0250 | 1 | 0.0123 | 0.32 | -0.0139 | 1 | -0.0369 | 1 | -0.0328 | 0.84 |
| Raw | spectral_sdm_raw_tissue | Tissue | -0.0100 | 1 | 0.0598 | 0 | -0.0056 | 0.8471 | 0.025 | 0.2824 | 0.0989 | 0 |
| Raw | spectral_landscape_raw_tissue | Tissue | 0.0881 | 0 | 0.1516 | 0 | 0.0515 | 0 | 0.1226 | 0 | 0.21378 | 0 |
| Raw | spectral_bdm_raw_sra | SRA | -0.0122 | 1 | 0.143 | 0 | -0.0123 | 1 | 0.0078 | 0.4 | -0.0928 | 1 |
| Raw | spectral_sdm_raw_sra | SRA | -0.0026 | 1 | 0.174 | 0 | -0.0126 | 1 | 0.0162 | 0.2824 | 0.0455 | 0.2667 |
| Raw | spectral_landscape_raw_sra | SRA | 0.1317 | 0 | 0.3134 | 0 | 0.0726 | 0 | 0.2332 | 0 | 0.2350 | 0 |
| Raw | spectral_bdm_raw_approach | Approach | -0.01284 | 1 | -0.0147 | 1 | 0.9309 | 0 | 0.9096 | 0 | 0.9852 | 0 |
| Raw | spectral_sdm_raw_approach | Approach | -0.0098 | 1 | -0.0038 | 1 | 0.9520 | 0 | 0.9330 | 0 | 0.9853 | 0 |
| Raw | spectral_landscape_raw_approach | Approach | -0.0277 | 1 | -0.0193 | 1 | 0.6937 | 0 | 0.7663 | 0 | 0.9848 | 0 |
| SCT Individual | seurat_sct_individual_sra | SRA | 0.0268 | 0 | 0.2020 | 0 | 0.0142 | 0 | 0.2020 | 0 | 0 | 1 |
| SCT Individual | seurat_sct_individual_tissue | Tissue | -0.0003 | 0.84 | 0.0524 | 0 | -0.0001 | 0.84 | 0.0524 | 0 | 0 | 1 |
| SCT Individual | seurat_sct_individual_approach | Approach | -0.1031 | 1 | -0.5218 | 1 | -0.0515 | 1 | -0.5218 | 1 | 0 | 1 |
| SCT Individual | kmeans_sct_individual_tissue | Tissue | 0.4568 | 0 | 0.4990 | 0 | 0.3259 | 0 | 0.5214 | 0 | 0.5630 | 0 |
| SCT Individual | kmeans_sct_individual_sra | SRA | 0.3515 | 0 | 0.5048 | 0 | 0.2228 | 0 | 0.4577 | 0 | 0.5977 | 0 |
| SCT Individual | kmeans_sct_individual_approach | Approach | -0.0165 | 1 | -0.0043 | 0.8727 | 0.9053 | 0 | 0.8894 | 0 | 0.9853 | 0 |
| SCT Individual | hierarchical_bdm_ph_sct_individual_tissue | Tissue | 0.0151 | 0 | 0.0469 | 0 | 0.0096 | 0 | 0.0089 | 0.2666 | 0.1029 | 0 |
| SCT Individual | hierarchical_sdm_ph_sct_individual_tissue | Tissue | 0.0033 | 0.2526 | 0.0068 | 0.2526 | 0.0025 | 0.1263 | -0.0601 | 1 | 0.0786 | 0 |
| SCT Individual | hierarchical_landscape_ph_sct_individual_tissue | Tissue | 0.1116 | 0 | 0.1937 | 0 | 0.0663 | 0 | 0.1595 | 0 | 0.2137 | 0 |
| SCT Individual | hierarchical_bdm_ph_sct_individual_sra | SRA | 0.0419 | 0 | 0.0582 | 0 | -0.0144 | 1 | -0.1172 | 1 | 0.0838 | 0 |
| SCT Individual | hierarchical_sdm_ph_sct_individual_sra | SRA | 0.0396 | 0 | 0.0535 | 0 | -0.0160 | 1 | -0.1317 | 1 | 0.0759 | 0 |
| SCT Individual | hierarchical_landscape_ph_sct_individual_sra | SRA | 0.1023 | 0 | 0.3066 | 0 | 0.0546 | 0 | 0.2294 | 0 | 0.1948 | 0 |
| SCT Individual | hierarchical_bdm_ph_sct_individual_approach | Approach | 0.1417 | 0 | 0.0370 | 0 | 0.9490 | 0 | 0.9234 | 0 | 0.9852 | 0 |
| SCT Individual | hierarchical_sdm_ph_sct_individual_approach | Approach | 0.1349 | 0 | 0.0190 | 0 | 0.9079 | 0 | 0.8829 | 0 | 0.9851 | 0 |
| SCT Individual | hierarchical_landscape_ph_sct_individual_approach | Approach | -0.0314 | 1 | -0.0112 | 1 | 0.4284 | 0 | 0.6803 | 0 | 0.9847 | 0 |
| SCT Individual | spectral_bdm_sct_individual_tissue | Tissue | 0.0437 | 0 | 0.0807 | 0 | 0.0270 | 0 | 0.0599 | 0 | 0.1349 | 0 |
| SCT Individual | spectral_sdm_sct_individual_tissue | Tissue | 0.0061 | 0.1333 | 0.0148 | 0 | 0.0042 | 0.1263 | -0.0453 | 1 | 0.0764 | 0 |
| SCT Individual | spectral_landscape_sct_individual_tissue | Tissue | 0.0682 | 0 | 0.1433 | 0 | 0.0394 | 0 | 0.1140 | 0 | 0.1545 | 0 |
| SCT Individual | spectral_bdm_sct_individual_sra | SRA | 0.0465 | 0 | 0.0892 | 0 | -0.0072 | 1 | -0.0470 | 1 | 0.0957 | 0 |
| SCT Individual | spectral_sdm_sct_individual_sra | SRA | 0.1403 | 0 | 0.2239 | 0 | 0.0693 | 0 | 0.1103 | 0 | 0.1434 | 0 |
| SCT Individual | spectral_landscape_sct_individual_sra | SRA | 0.0697 | 0 | 0.2582 | 0 | 0.0362 | 0 | 0.1758 | 0 | 0.1116 | 0 |
| SCT Individual | spectral_bdm_sct_individual_approach | Approach | 0.2887 | 0 | 0.1422 | 0 | 0.9770 | 0 | 0.9644 | 0 | 0.9879 | 0 |
| SCT Individual | spectral_sdm_sct_individual_approach | Approach | 0.0385 | 0 | -0.0053 | 0.8727 | 0.8891 | 0 | 0.8722 | 0 | 0.9851 | 0 |
| SCT Individual | spectral_landscape_sct_individual_approach | Approach | -0.0279 | 1 | -0.0085 | 0.8727 | 0.4058 | 0 | 0.6751 | 0 | 0.9847 | 0 |
| SCT Whole | seurat_sct_whole_sra | SRA | 0.2503 | 0 | 0.4686 | 0 | 0.1484 | 0 | 0.5557 | 0 | 0.8560 | 0 |
| SCT Whole | seurat_sct_whole_tissue | Tissue | 0.2131 | 0 | 0.3786 | 0 | 0.1247 | 0 | 0.5232 | 0 | 0.9522 | 0 |
| SCT Whole | seurat_sct_whole_approach | Approach | -0.0229 | 1 | -0.0514 | 1 | -0.0077 | 1 | 0.1920 | 0 | 0.8896 | 0 |
| SCT Whole | kmeans_sct_whole_tissue | Tissue | 0.4668 | 0 | 0.5308 | 0 | 0.3322 | 0 | 0.5231 | 0 | 0.5675 | 0 |
| SCT Whole | kmeans_sct_whole_sra | SRA | 0.4109 | 0 | 0.5454 | 0 | 0.2706 | 0 | 0.5082 | 0 | 0.6121 | 0 |
|  |  |  | **ARI** |  | **NMI** |  | **Jaccard** |  | **VI** |  | **Purity** |  |
| **Iteration** | **Method** | **ReferenceCol** | **Norm** | **FDR** | **Norm** | **FDR** | **Norm** | **FDR** | **Norm** | **FDR** | **Norm** | **FDR** |
| SCT Whole | kmeans_sct_whole_approach | Approach | 0.1427 | 0 | 0.0451 | 0 | 0.6900 | 0 | 0.7061 | 0 | 0.8162 | 0 |
| SCT Whole | hierarchical_bdm_ph_sct_whole_tissue | Tissue | 0.1111 | 0 | 0.0842 | 0 | 0.0687 | 0 | 0.0568 | 0 | 0.1050 | 0 |
| SCT Whole | hierarchical_sdm_ph_sct_whole_tissue | Tissue | -0.0115 | 1 | 0.0200 | 0 | -0.0063 | 0.9818 | -0.0267 | 1 | 0.0635 | 0.12 |
| SCT Whole | hierarchical_landscape_ph_sct_whole_tissue | Tissue | 0.0612 | 0 | 0.1168 | 0 | 0.0355 | 0 | 0.0778 | 0 | 0.1424 | 0 |
| SCT Whole | hierarchical_bdm_ph_sct_whole_sra | SRA | 0.0557 | 0 | 0.2004 | 0 | 0.0201 | 0 | 0.0468 | 0 | 0.0179 | 0.2181 |
| SCT Whole | hierarchical_sdm_ph_sct_whole_sra | SRA | -0.0182 | 1 | 0.0665 | 0 | -0.0363 | 1 | -0.0603 | 1 | 0.0429 | 0.2181 |
| SCT Whole | hierarchical_landscape_ph_sct_whole_sra | SRA | 0.0836 | 0 | 0.2673 | 0 | 0.0441 | 0 | 0.1833 | 0 | 0.1535 | 0 |
| SCT Whole | hierarchical_bdm_ph_sct_whole_approach | Approach | 0.0983 | 0 | 0.0042 | 0.24 | 0.6359 | 0 | 0.6444 | 0 | 0.8001 | 0 |
| SCT Whole | hierarchical_sdm_ph_sct_whole_approach | Approach | -0.0336 | 1 | -0.0104 | 1 | 0.6420 | 0 | 0.6845 | 0 | 0.7997 | 0 |
| SCT Whole | hierarchical_landscape_ph_sct_whole_approach | Approach | -0.0044 | 0.8842 | -0.0073 | 0.7636 | 0.3130 | 0 | 0.5095 | 0 | 0.7959 | 0 |
| SCT Whole | spectral_bdm_sct_whole_tissue | Tissue | 0.0643 | 0 | 0.0972 | 0 | 0.0377 | 0 | 0.0548 | 0 | 0.1627 | 0 |
| SCT Whole | spectral_sdm_sct_whole_tissue | Tissue | -0.0077 | 0.8842 | 0.0290 | 0.1333 | -0.0042 | 0.6857 | -0.0106 | 0.6545 | 0.0405 | 0.12 |
| SCT Whole | spectral_landscape_sct_whole_tissue | Tissue | 0.0365 | 0.15 | 0.1105 | 0 | 0.0209 | 0.12 | 0.0757 | 0 | 0.0981 | 0 |
| SCT Whole | spectral_bdm_sct_whole_sra | SRA | 0.0257 | 0 | 0.2020 | 0 | 0.0079 | 0 | 0.0807 | 0 | 0.0231 | 0.3 |
| SCT Whole | spectral_sdm_sct_whole_sra | SRA | 0.0649 | 0 | 0.2147 | 0 | 0.0319 | 0 | 0.1024 | 0 | 0.0431 | 0.12 |
| SCT Whole | spectral_landscape_sct_whole_sra | SRA | 0.0431 | 0 | 0.2284 | 0 | 0.0210 | 0 | 0.1366 | 0 | 0.0396 | 0.3 |
| SCT Whole | spectral_bdm_sct_whole_approach | Approach | 0.0983 | 0 | 0.0042 | 0.24 | 0.6359 | 0 | 0.6444 | 0 | 0.8001 | 0 |
| SCT Whole | spectral_sdm_sct_whole_approach | Approach | -0.0796 | 1 | 0.0145 | 0.1333 | 0.5646 | 0 | 0.6409 | 0 | 0.8003 | 0 |
| SCT Whole | spectral_landscape_sct_whole_approach | Approach | -0.0089 | 0.8842 | -0.0081 | 0.7636 | 0.3086 | 0 | 0.5085 | 0 | 0.7966 | 0 |
| Integrated | seurat_integrated_sra | SRA | 0.0726 | 0 | 0.1378 | 0 | 0.0362 | 0 | 0.0386 | 0 | 0.2903 | 0 |
| Integrated | seurat_integrated_tissue | Tissue | 0.0699 | 0 | 0.1190 | 0 | 0.0384 | 0 | 0.1169 | 0 | 0.3185 | 0 |
| Integrated | seurat_integrated_approach | Approach | -0.0072 | 1 | -0.0193 | 1 | 0.0143 | 0 | 0.3322 | 0 | 0.9845 | 0 |
| Integrated | kmeans_integrated_tissue | Tissue | 0.0115 | 0 | 0.0362 | 0 | 0.0066 | 0 | -0.0084 | 1 | 0.1407 | 0 |
| Integrated | kmeans_integrated_sra | SRA | 0.0275 | 0 | 0.0982 | 0 | 0.0089 | 0 | -0.0281 | 1 | 0.2122 | 0 |
| Integrated | kmeans_integrated_approach | Approach | -0.0042 | 1 | -0.0009 | 1 | 0.5705 | 0 | 0.7247 | 0 | 0.9854 | 0 |
| Integrated | hierarchical_bdm_ph_integrated_tissue | Tissue | 0.0203 | 0 | -0.0090 | 1 | 0.0125 | 0 | -0.0689 | 1 | 0.0372 | 0.1142 |
| Integrated | hierarchical_sdm_ph_integrated_tissue | Tissue | 0.0221 | 0 | 0.0294 | 0 | 0.0138 | 0 | -0.0179 | 1 | 0.0823 | 0 |
| Integrated | hierarchical_landscape_ph_integrated_tissue | Tissue | 0.1886 | 0 | 0.2533 | 0 | 0.1164 | 0 | 0.2232 | 0 | 0.2749 | 0 |
| Integrated | hierarchical_bdm_ph_integrated_sra | SRA | 0.0309 | 0 | 0.1362 | 0 | 0.0003 | 0.4173 | 0.0082 | 0.3 | 0.0191 | 0.3 |
| Integrated | hierarchical_sdm_ph_integrated_sra | SRA | 0.0427 | 0 | 0.0609 | 0 | -0.0092 | 1 | -0.0875 | 1 | 0.0483 | 0.2181 |
| Integrated | hierarchical_landscape_ph_integrated_sra | SRA | 0.1383 | 0 | 0.3514 | 0 | 0.0765 | 0 | 0.2769 | 0 | 0.2126 | 0 |
| Integrated | hierarchical_bdm_ph_integrated_approach | Approach | -0.0196 | 1 | -0.0110 | 1 | 0.8425 | 0 | 0.8440 | 0 | 0.9852 | 0 |
| Integrated | hierarchical_sdm_ph_integrated_approach | Approach | -0.0110 | 1 | -0.0087 | 1 | 0.9468 | 0 | 0.9266 | 0 | 0.9852 | 0 |
| Integrated | hierarchical_landscape_ph_integrated_approach | Approach | -0.0261 | 1 | -0.0143 | 1 | 0.6849 | 0 | 0.7644 | 0 | 0.9848 | 0 |
| Integrated | spectral_bdm_integrated_tissue | Tissue | 0.0034 | 0.45 | 0.0445 | 0 | 0.0020 | 0.3272 | -0.0009 | 0.6666 | 0.0176 | 0.3 |
| Integrated | spectral_sdm_integrated_tissue | Tissue | 0.0101 | 0.32 | 0.0227 | 0.32 | 0.0064 | 0 | -0.0222 | 1 | 0.1144 | 0 |
| Integrated | spectral_landscape_integrated_tissue | Tissue | 0.1579 | 0 | 0.2357 | 0 | 0.0955 | 0 | 0.2134 | 0 | 0.2705 | 0 |
| Integrated | spectral_bdm_integrated_sra | SRA | 0.0450 | 0 | 0.2084 | 0 | 0.0196 | 0 | 0.0907 | 0 | 0.0474 | 0 |
| Integrated | spectral_sdm_integrated_sra | SRA | 0.0470 | 0 | 0.1105 | 0 | 0.0037 | 0.3272 | -0.0056 | 0.6666 | 0.0578 | 0.1142 |
| Integrated | spectral_landscape_integrated_sra | SRA | 0.1149 | 0 | 0.3023 | 0 | 0.0628 | 0 | 0.2309 | 0 | 0.1856 | 0 |
| Integrated | spectral_bdm_integrated_approach | Approach | -0.0110 | 1 | -0.0087 | 1 | 0.9468 | 0 | 0.9266 | 0 | 0.9852 | 0 |
| Integrated | spectral_sdm_integrated_approach | Approach | -0.0105 | 1 | -0.0080 | 1 | 0.9495 | 0 | 0.9296 | 0 | 0.9853 | 0 |
| Integrated | spectral_landscape_integrated_approach | Approach | -0.0330 | 1 | -0.0126 | 1 | 0.4432 | 0 | 0.6839 | 0 | 0.9846 | 0 |

##### **Supplemental Table 6: Within-Iteration Results, Validation Dataset** (file: Supplemental_Table_6.csv)

**Supplemental Table 7: Cross-Iteration Results, Validation Dataset** (file: Supplemental_Table_7.csv)

**Supplemental Table 8: Within-Iteration Results, Main Dataset** (file: Supplemental_Table_8.csv)

**Supplemental Table 9: Cross-Iteration Results, Main Dataset** (file: Supplemental_Table_9.csv)

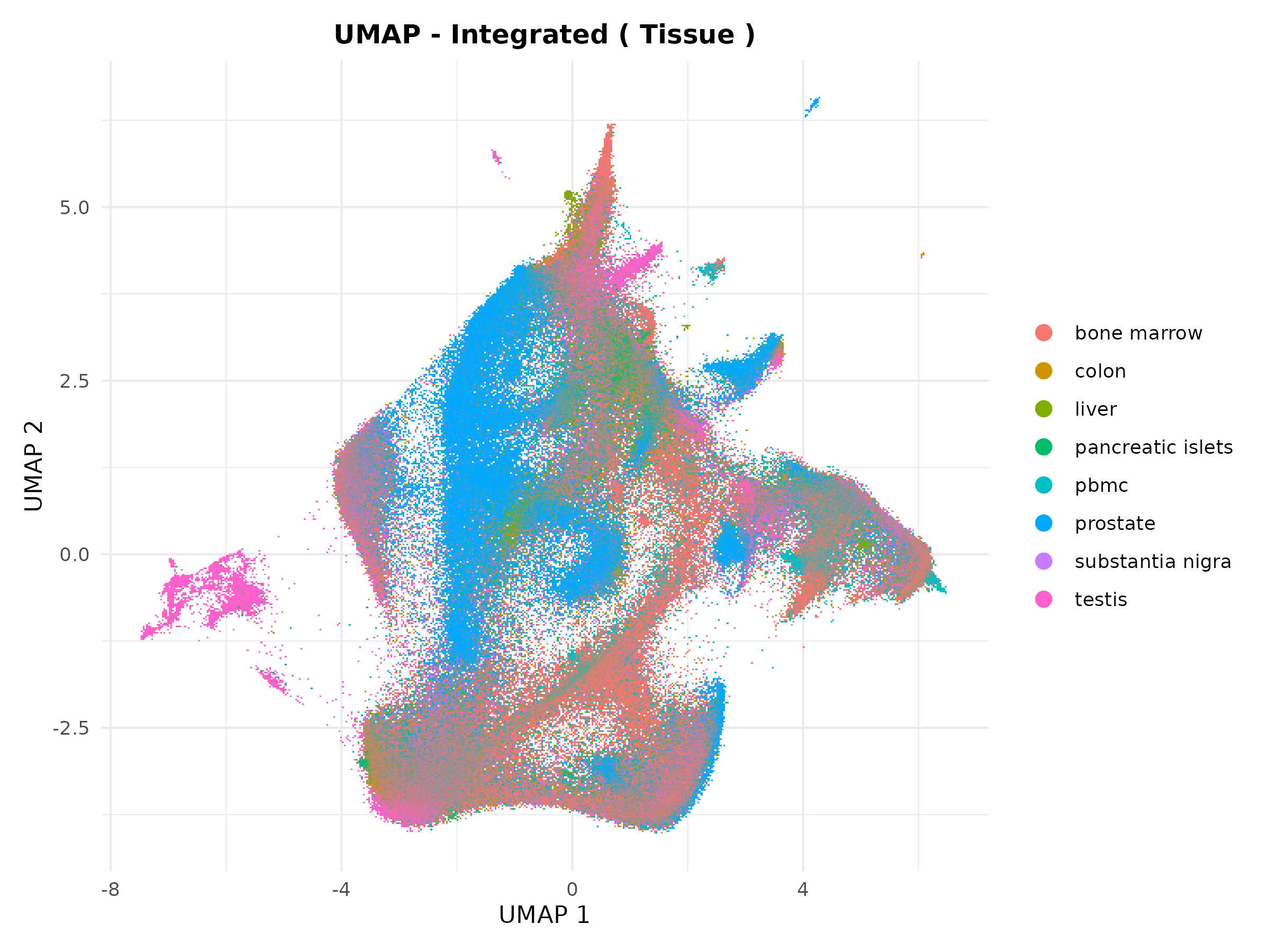

**Supplemental Figure 1A: Integrated UMAP labelled by Tissue.**

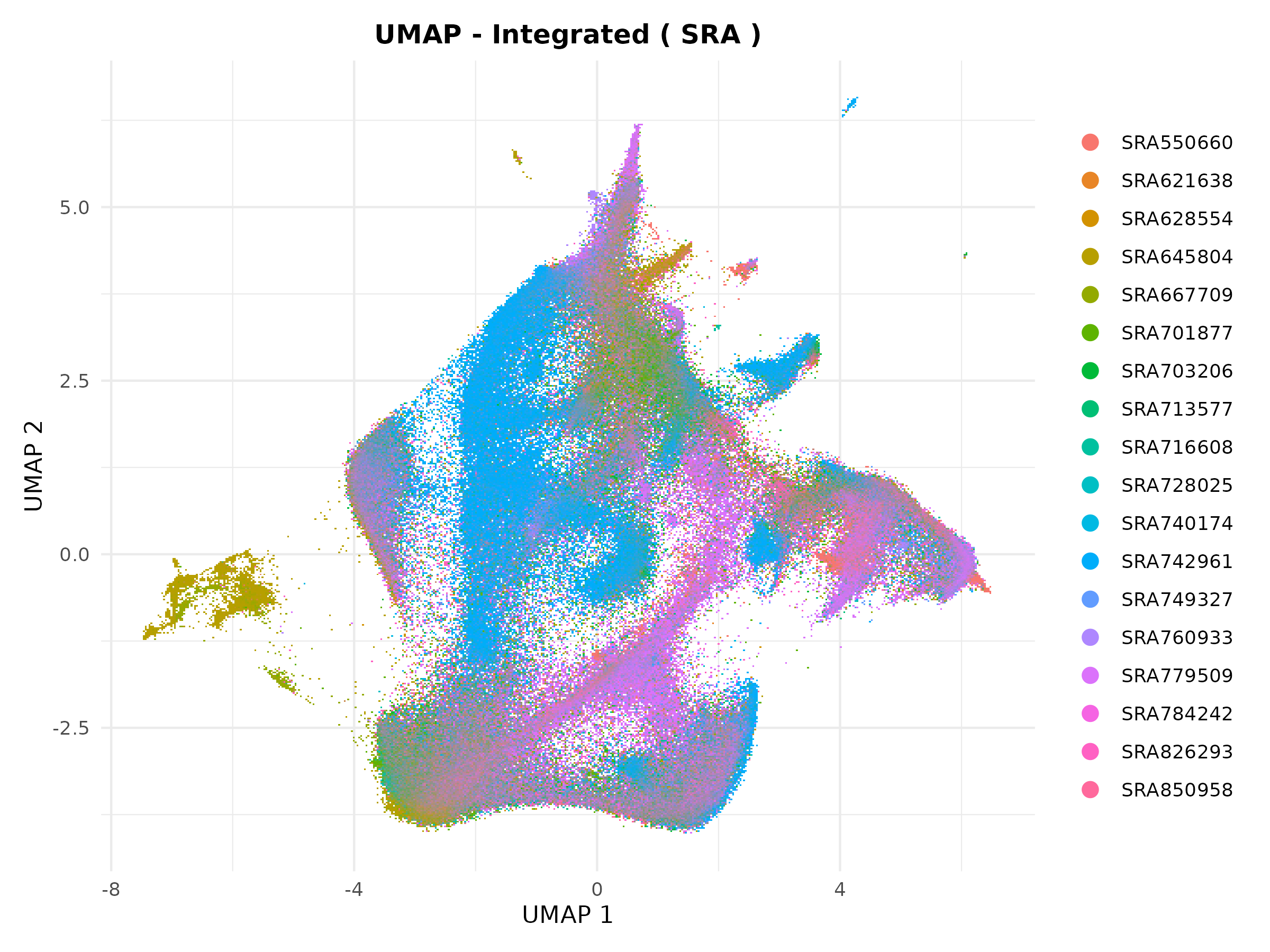

**Supplemental Figure 1B: Integrated UMAP labelled by SRA/Study Origin.**

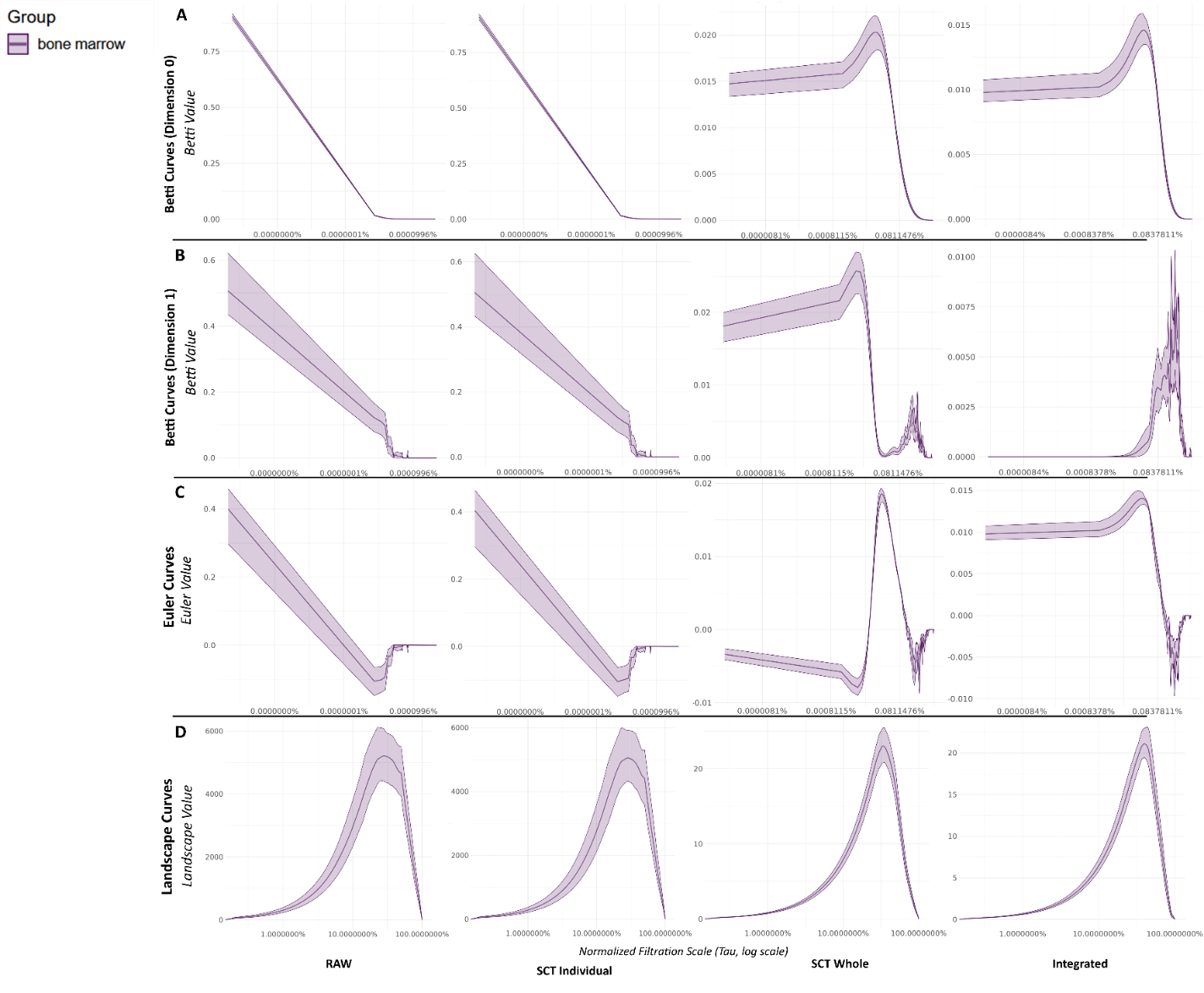

**Supplemental Figure 2: Mean Betti, Euler, and Landscape Curves for Bone Marrow Data (GSE120221).** The figure displays the mean topological summary curves for the entire GSE120221 bone marrow dataset, illustrating the average topological signature of this homogeneous cell population across different preprocessing workflows. Each column represents a distinct data preprocessing iteration: Raw, SCT Individual, SCT Whole, and Integrated. The rows show different topological summaries: (A) Betti 0 (β0), (B) Betti 1 (β1), (C) Euler characteristic (χ=β0−β1), and (D) aggregated Landscape curves. For each plot, the solid line represents the mean curve calculated across all 25 samples, and the shaded ribbon indicates the 95% confidence interval derived from bootstrapping. These plots demonstrate how the average topological profile and its within-group variability change as a result of normalization and integration.

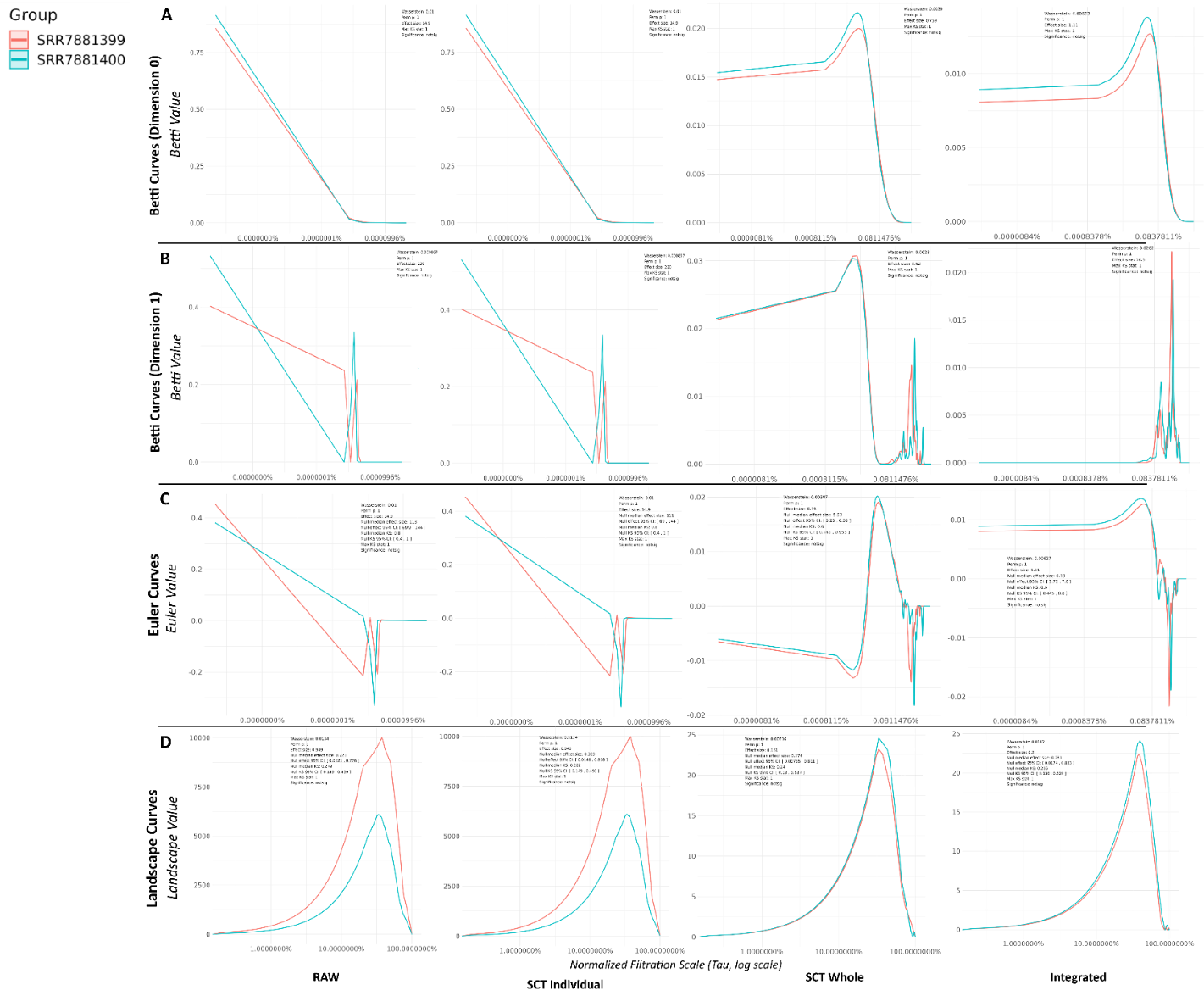

**Supplemental Figure 3: Pairwise Comparison of Betti, Euler, and Landscape Curves for Samples SRR7881399 and SRR7881400.** This figure displays Betti 0 (β_0_) (A), Betti 1 (β_1_) (B), Euler (C), and Landscape (D) curves for two samples (SRR7881399, salmon; SRR7881400, teal) across four processing stages shown in columns: RAW, SCT individual, SCT Whole, and Integrated. While initially distinct, the curves for the two samples become progressively more aligned with each step. The final near-perfect overlap in the Integrated column demonstrates that the workflow successfully homogenizes the data and corrects for batch effects. Full statistics are in supplemental Table 6.

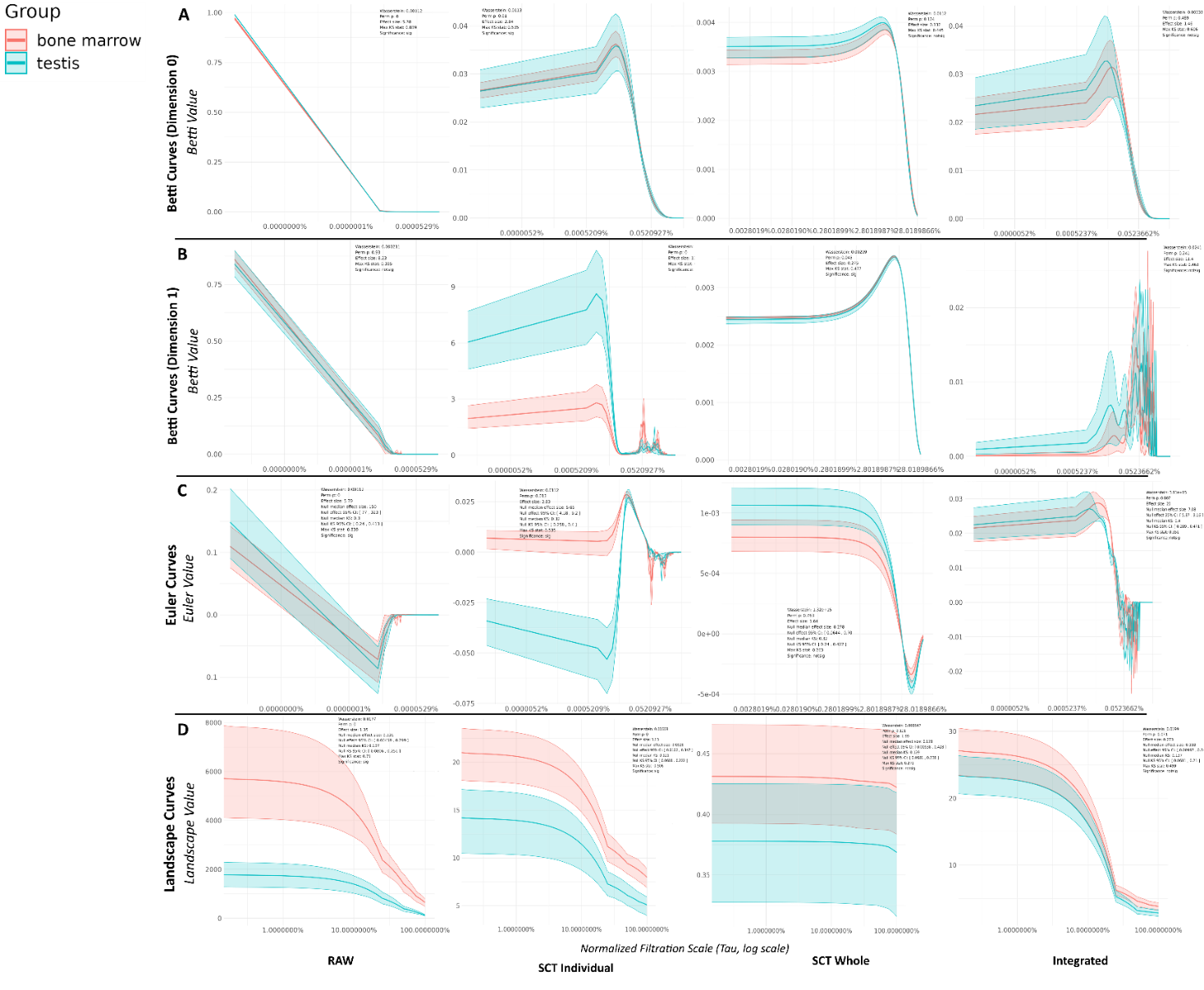

**Supplemental Figure 4: Cross-Iteration Comparison of Mean Topological Summary Curves for Bone Marrow vs. Testis.** This figure compares the mean topological curves (with 95% confidence intervals) for bone marrow (teal) and testis (salmon) tissues. The rows display different topological statistics, Betti 0 (β0) (A), Betti 1 (β1) (B), Euler (C), and Landscape (D, across the four data processing stages shown in the columns (RAW to Integrated). The distinct topological signatures of these biologically different tissues remain clearly separated throughout all processing steps, demonstrating that the integration pipeline preserves true biological variation while correcting for technical noise. Full statistics are in Supplemental Table 8.
